# Comprehensive mapping of SARS-CoV-2 interactions in vivo reveals functional virus-host interactions

**DOI:** 10.1101/2021.01.17.427000

**Authors:** Siwy Ling Yang, Louis DeFalco, Danielle E. Anderson, Yu Zhang, Ashley J Aw, Su Ying Lim, Lim Xin Ni, Kiat Yee Tan, Tong Zhang, Tanu Chawla, Yan Su, Alexander Lezhava, Paola de Sessions, Andres Merits, Lin-Fa Wang, Roland G. Huber, Yue Wan

**Affiliations:** Epigenetic and Epitranscriptomic Regulation, Genome Institute of Singapore, Agency for Science, Technology and Research (A*STAR), Singapore 138672, Singapore; Biomolecular Function Discovery, Bioinformatics Institute (BII), Agency for Science, Technology and Research (A*STAR), Matrix #07-01, Singapore 138671; Programme in Emerging Infectious Diseases, Duke-NUS Medical School, 8 College Road, 169857, Singapore; Laboratory of translational diagnostics, Genome Institute of Singapore, Agency for Science, Technology and Research (A*STAR), Singapore 138672, Singapore; Oxford Nanopore Technologies; Institute of Technology, University of Tartu, Tartu, Estonia; School of Biological Sciences, Nanyang Technological University, Singapore 637551; Department of Biochemistry, Yong Loo Lin School of Medicine, National University of Singapore, Singapore 117597, Singapore

**Keywords:** SARS-CoV-2, COVID-19, RNA structure probing, structure modelling, high throughput sequencing, proximity ligation, SHAPE, interactome, nanopore, epitranscriptomics

## Abstract

SARS-CoV-2 has emerged as a major threat to global public health, resulting in global societal and economic disruptions. Here, we investigate the intramolecular and intermolecular RNA interactions of wildtype (WT) and a mutant (Δ382) SARS-CoV-2 virus in cells using high throughput structure probing on Illumina and Nanopore platforms. We identified twelve potentially functional structural elements within the SARS-CoV-2 genome, observed that identical sequences can fold into divergent structures on different subgenomic RNAs, and that WT and Δ382 virus genomes can fold differently. Proximity ligation sequencing experiments identified hundreds of intramolecular and intermolecular pair-wise interactions within the virus genome and between virus and host RNAs. SARS-CoV-2 binds strongly to mitochondrial and small nucleolar RNAs and is extensively 2’-O-methylated. 2’-O-methylation sites in the virus genome are enriched in the untranslated regions and are associated with increased pair-wise interactions. SARS-CoV-2 infection results in a global decrease of 2’-O-methylation sites on host mRNAs, suggesting that binding to snoRNAs could be a pro-viral mechanism to sequester methylation machinery from host RNAs towards the virus genome. Collectively, these studies deepen our understanding of the molecular basis of SARS-CoV-2 pathogenicity, cellular factors important during infection and provide a platform for targeted therapy.

## Introduction

Coronaviruses (CoVs) are enveloped viruses with positive-sense single-stranded RNA genomes. They are widespread in animals and can cause mild to severe respiratory or enteric disease in humans^1,2^. There are currently seven CoVs known to infect humans, which include the four “seasonal” human CoVs: OC43, 229E, NL63 and HKU1^3^, which can cause mild cold-like symptoms and highly pathogenic CoVs: SARS-CoV, MERS-CoV, and SARS-CoV-2. SARS-CoV emerged in 2002 and resulted in more than 8,000 human infections with case fatality rate of approximately 10%^4^. This was followed by the discovery of MERS-CoV in 2012, which has resulted in more than 2,000 human infections and over 800 lethal cases with ongoing sporadic outbreaks in the Middle East^5^. More recently, the outbreak of SARS-CoV-2 has caused unprecedented social and economic damage around the world^6^. Since it was first reported in Wuhan, China in December 2019, SARS-CoV-2 has resulted in over 90 million infections and more than 1.9 million deaths as of 13^th^ January 2021, according to WHO. Different variants of SARS-CoV-2 have been found to circulate within patients. Viruses that contain deletions of various sizes in the ORF8 region has been found around the world, including in Singapore, Taiwan, Bangladesh, Australia and Spain^7^. In particular, a 382-nucleotide deletion (Δ382) of the SARS-CoV-2 genome that truncates ORF7 and deletes ORF8 was found in patients in Singapore^8^. While patients infected with Δ382 virus showed less severe symptoms than those infected with WT viruses, the molecular mechanisms behind virus attenuation in patients is unclear^7^. It is hence imperative to understand how SARS-CoV-2 and their variants function, in order to facilitate effective surveillance, prevention and treatment strategies.

CoV genomes are among the largest of the RNA viruses, with lengths of 26-32 kb^2^. Upon entry into the cell, the positive-sense genome is translated from two open reading frames (ORF1a and ORF1b) and the resulting polyproteins are cleaved into non-structural proteins. Non-structural proteins are essential for virus RNA replication that, in addition to new genomes, also generates numerous sub-genomic RNA (sgRNA) species using discontinuous transcription^9^. These RNAs, together with the full genome, can interact with host cell proteins and RNAs to regulate virus infection. Like many other RNA viruses such as dengue virus (DENV) and Zika virus (ZIKV), the SARS-CoV-2 genomic RNA can fold into secondary and tertiary structures that are essential for virus RNA replication and protein translation^10–13^. Importantly, elements in the 5’ and 3’ untranslated regions (UTRs) have been implicated in virus replication and protein synthesis^14,15^, and the frameshifting element is important for ribosome slippage to enable the translation of ORF1ab^16,17^. However, how the rest of the virus genomes folds into short- and long-range structures is still under-studied. Here, we utilized different high throughput RNA and interactome techniques (SHAPE-MaP^18^, PORE-cupine^19^ and SPLASH^20^) to comprehensively interrogate the secondary structures and virus-host interactions along the wildtype (WT) and Δ382 SARS-CoV-2 genomes to identify potentially functional structure elements along the virus genome (**Figure 1a**). Using PORE-cupine, we identified sgRNA-specific structures as well as WT and Δ382 specific structures using Nanopore direct RNA sequencing. The advantage of long-read sequencing enables us to map our sequencing reads uniquely to each sgRNA, without needing to average structure signals across all sgRNAs and full-length RNA in short-read sequencing, due to ambiguous mapping. SPLASH further allows us to identify pair-wise RNA interactions using proximity ligation and sequencing, deepening our knowledge of how the genome folds along itself for function.

**Figure 1.**
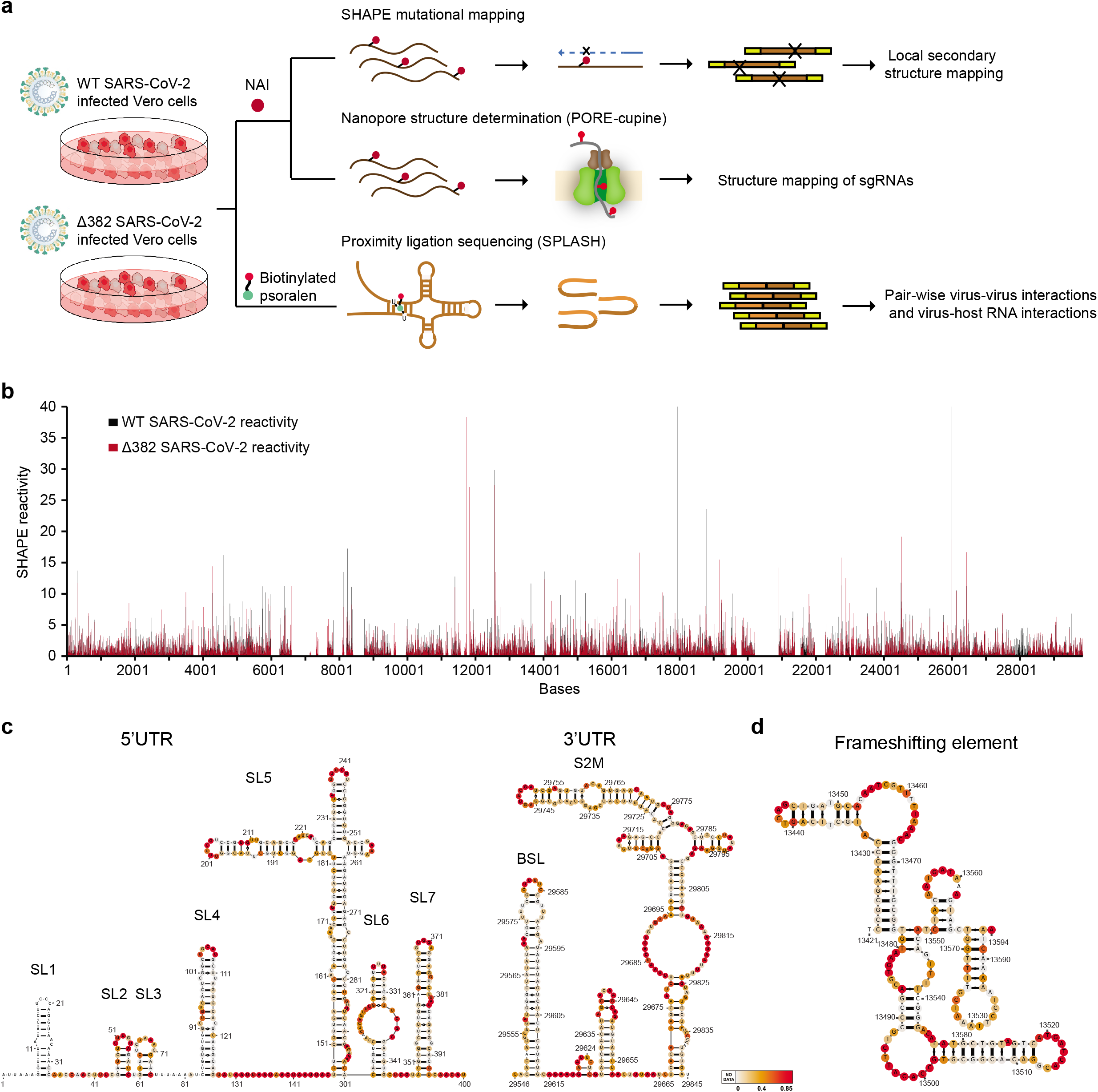
Comprehensive structure and interactome mapping of SARS-CoV-2 RNA in cell. **a**, Schematic of the different strategies that were used to probe WT and Δ382 inside infected Vero cells. NAI was used to modify single-stranded bases in cells and these modifications are then either read out directly using Nanopore direct RNA sequencing or converted into a cDNA library for Illumina sequencing. Biotinylated psoralen was used to crosslink pair-wise RNA interactions in vivo to capture both intramolecular and intermolecular RNA-RNA interactions. **b**, SHAPE-reactivity along the WT (black) and Δ382 (red) SARS-CoV-2 genome. Higher reactivity regions tend to be more single-stranded. **c**,**d**, Structure models are generated using the program RNA Structure using SHAPE-reactivities as constraints^53^, and visualized using VARNA^56^. The modelled structures of 5’UTR and 3’UTR (**c**), and that of the frameshift element (**d**) agree with known models in the literature.

In addition to studying how the virus genome interacts with itself, determining how the virus genome interacts with host RNAs in its cellular environment is another key to understanding virus pathogenicity. Other RNA viruses, including the ZIKV, have been shown to directly interact with host RNAs such as microRNAs to impact virus infection^11^. Most host factor studies for SARS-CoV-2 done to date have been focused on how the host proteins interact with the virus proteins and genome; much less is known about how SARS-CoV-2 interacts with host RNAs inside cells^21–24^. Here, we utilized SPLASH to identify host RNAs that interact with the SARS-CoV-2 genome inside infected Vero cells (**Figure 1a**). We observed that SARS-CoV-2 RNA interacts strongly with a small nucleolar RNA (snoRNA) SNORD27 and is 2’-O-methylated inside cells. We further showed that 2’-O-methylation of host RNAs is decreased in SARS-CoV-2 infected cells, and that virus-SNORD27 interaction could serve as a mechanism for SARS-CoV-2 to facilitate host RNA degradation.

## Results

### SARS-CoV-2 RNA is highly structured in host cells

To study the secondary structures of SARS-CoV-2 RNAs inside cells, we infected Vero cells with WT and Δ382 SARS-CoV-2 and performed structure probing using the compound NAI inside cells (**Methods**)^10,25^. We then performed mutational mapping (MaP) to determine the location of high reactivity bases, indicating single-stranded bases, along the virus genome. We confirmed that mutational rates along NAI-treated, denatured and DMSO-treated samples are as expected (**Supp. Figure 1a**,**b, Supp. Table 1**). Biological replicates of SARS-CoV-2 SHAPE-MaP show that the reactivities across replicates are highly reproducible (**Supp. Figure 1c**), cover around 80% of the entire SARS-CoV-2 genome (**Figure 1b**), and map to known structures in the 5’ and 3’UTR as expected (**Supp. Figure 1d, Supp. Table 2**). SHAPE-MaP reactivities were used to constrain RNA secondary structure predictions to obtain accurate structure models of the entire SARS-CoV-2 genome (**Methods, Figure 1c**,**d, Figure 2**)^26^. Our structure models are consistent with previously identified structural elements in the 5’ and 3’UTRs (**Figure 1c**)^27–29^, existing structure models for the frameshifting element (**Figure 1d, Supp. Figure 2a**) and that for TRS-L elements (**Supp. Figure 2b**)^27^, confirming that models based on our data are accurate. Similar to other RNA viruses like DENV and ZIKV, 57% of the bases in the SARS-CoV-2 genome are predicted to be paired, with a median helix length of 5 bases in both WT and Δ382 genomes (**Supp. Figure 2c**)^10^. These short helices enable RNA viruses to escape from host immune responses.

**Figure 2.**
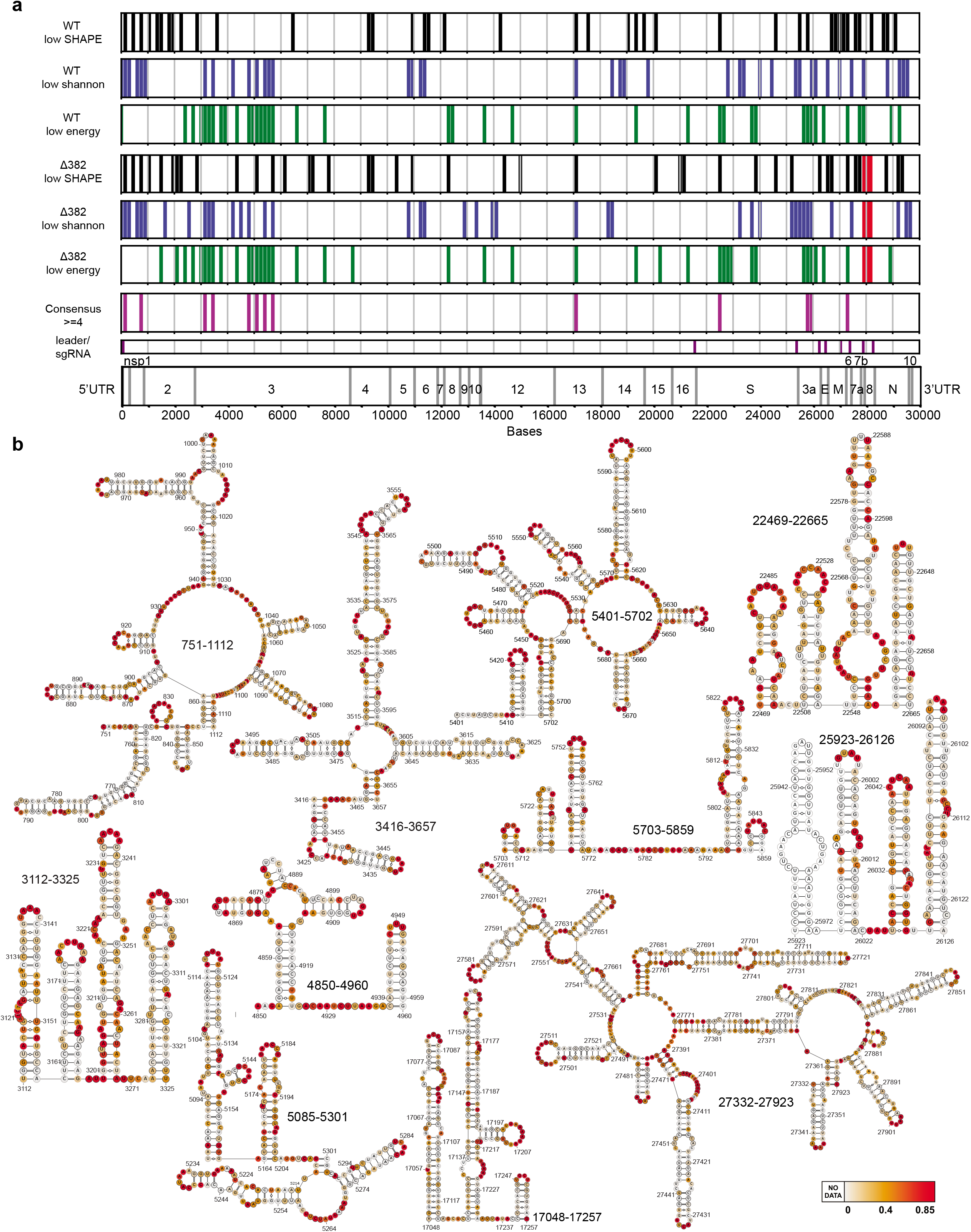
SHAPE-MaP identifies functional structural elements along the SARS-CoV-2 genome. **a**, Regions (150 bases) with the lowest 20% SHAPE reactivity (in black), Shannon entropy (blue) and greatest difference (top 20%) between actual and shuffled energies (green) are shown along both the WT and Δ382 genomes. 12 consensus regions are consistently highlighted (4/6) across both WT and Δ382 genomes and are shown in purple. The red box indicates the deletion region in the Δ382 genome. **b**, Structure models of the 12 consensus were generated using the program RNA Structure^53^, using SHAPE-MaP reactivities as constraints, and visualized using VARNA^56^. The SHAPE-reactivities are mapped onto the structure models.

To identify potentially functional structural RNA elements in the SARS-CoV-2 genome, we used a consensus model between WT and Δ382, incorporating local SHAPE reactivity, local Shannon entropy of the structure models and local =Scan-Fold’ energies in 150 nt windows (**Figure 2a**)^30,31^. We evaluated window sizes of 50 nt to 300 nt which yielded consistent results (**Supp. Figure 3a**). We considered a position a consensus candidate if it had at least four out of six possible characteristics of =low average SHAPE’ as an indication of structuredness at a location, =low Shannon entropy’ as an indication of structural consistency and limited alternative folding, and =low Scan-Fold energy’ as a proxy for high stability of putative structural elements in both WT and Δ382 (**Figure 2a**). Local structure models of consensus regions were largely consistent with structures obtained in the global context and identified novel highly structured elements within the genome (**Figure 2b**). As single-stranded regions present in the SARS-CoV-2 genome could be used for siRNA targeting, we also identified locations with high reactivities (top 20%) in both the WT and Δ382 genomes (**Supp. Figure 3b**). We identified a total of 21 regions that could be used for siRNA targeting, to facilitate potential treatments for COVID-19 disease.

**Figure 3.**
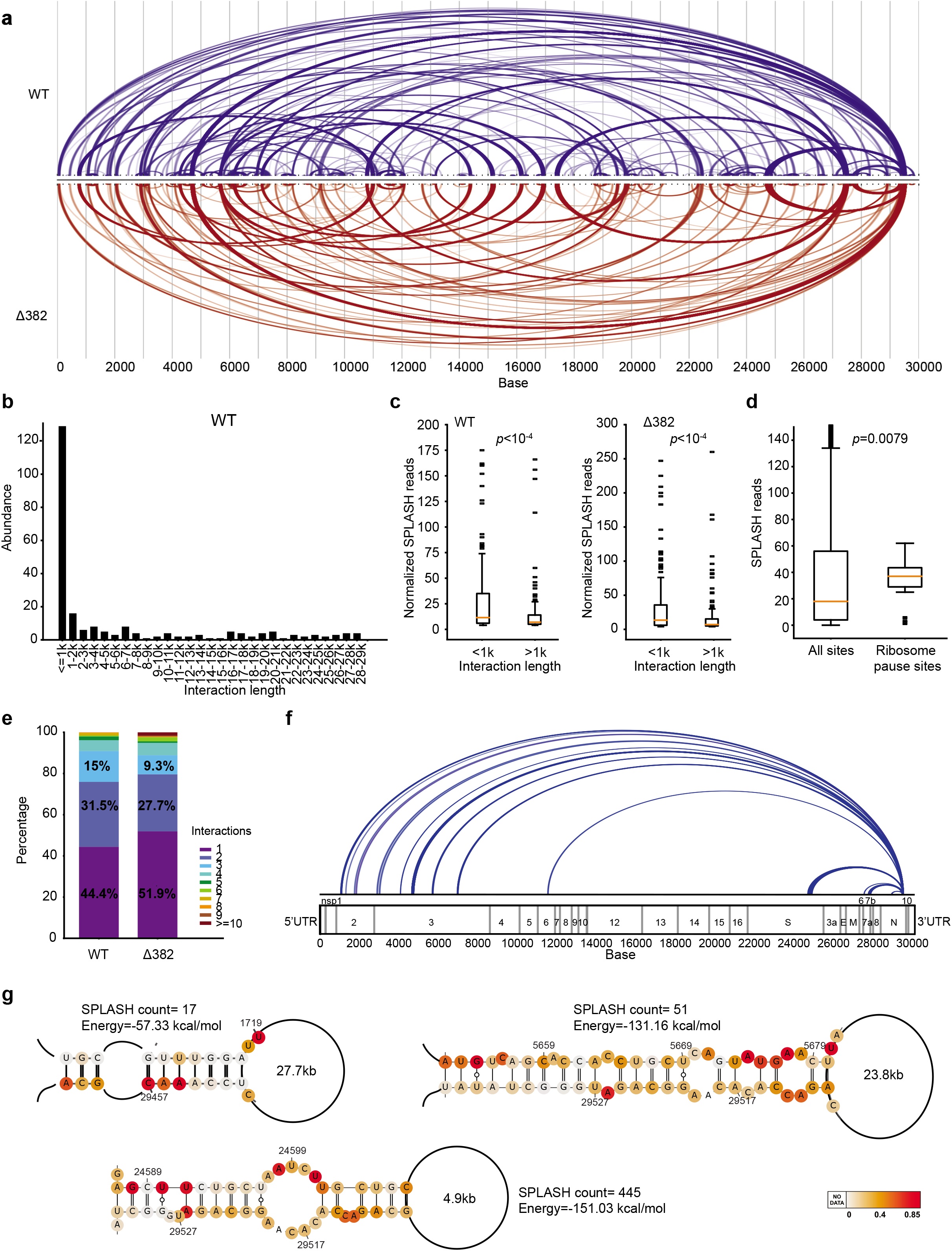
SARS-CoV-2 contains hundreds of intramolecular long-range interactions. **a**, Pair-wise RNA-RNA interactions along the WT (blue) and Δ382 (red) genomes. The thickness of the lines indicate the abundance of chimeric reads for that particular interaction. **b**, Histogram showing the distribution of interactions that span different lengths along the WT SARS-CoV-2 genome. Interactions longer than 1kb are classified as “long-range” and comprise 45.6% of all interactions. **c**, Boxplot showing the distribution of the abundance of SPLASH chimeric reads for long (>1kb) and short (<=1kb) pair-wise interactions in both WT (left) and Δ382 (right) genomes. Long interactions tend to have lower SPLASH interaction counts, suggesting that they may be formed more transiently. *P*-value is calculated using Wilcoxon Rank Sum test. **d**, Boxplot showing the distribution of SPLASH chimeric reads along the SARS-CoV-2 genome for all sites (left) and for sites that show ribosome pausing events (right). Sites with ribosome pausing events show higher SPLASH chimeric reads, indicating that they reside in more highly structured regions. *P*-value is calculated using Wilcoxon Rank Sum test. **e**, Barcharts showing the proportion of unique pair-wise interactions, as well as interactions that have 2 or more alternative partners, along the WT and Δ382 genomes. **f**, Arc plots showing the alternative interactions between N and other positions along the SARS-CoV-2 genome. **g**, Representative structure models for interactions between N and other regions along the genome. Structure models are generated using RNAcofold from regions identified by SPLASH. For each interaction, the SPLASH count and predicted energy of folding from RNAcofold is shown next to the model^34^. SHAPE-MaP reactivities are mapped onto the bases in the structure models.

### SARS-CoV-2 genome contains hundreds of regions involved in intramolecular long-range interactions

In addition to determining which bases are paired or unpaired in the SARS-CoV-2 genome, we also wanted to know the identity of pairing partners within the genome. To identify pair-wise RNA interactions, we treated SARS-CoV-2 or Δ382 infected Vero-E6 cells with biotinylated psoralen and performed proximity ligation sequencing (SPLASH)^20^. Biological replicates of SPLASH showed good correlation of pair-wise interactions between the samples, indicating that our method is robust (**Supp. Figure 4a,b**).

**Figure 4.**
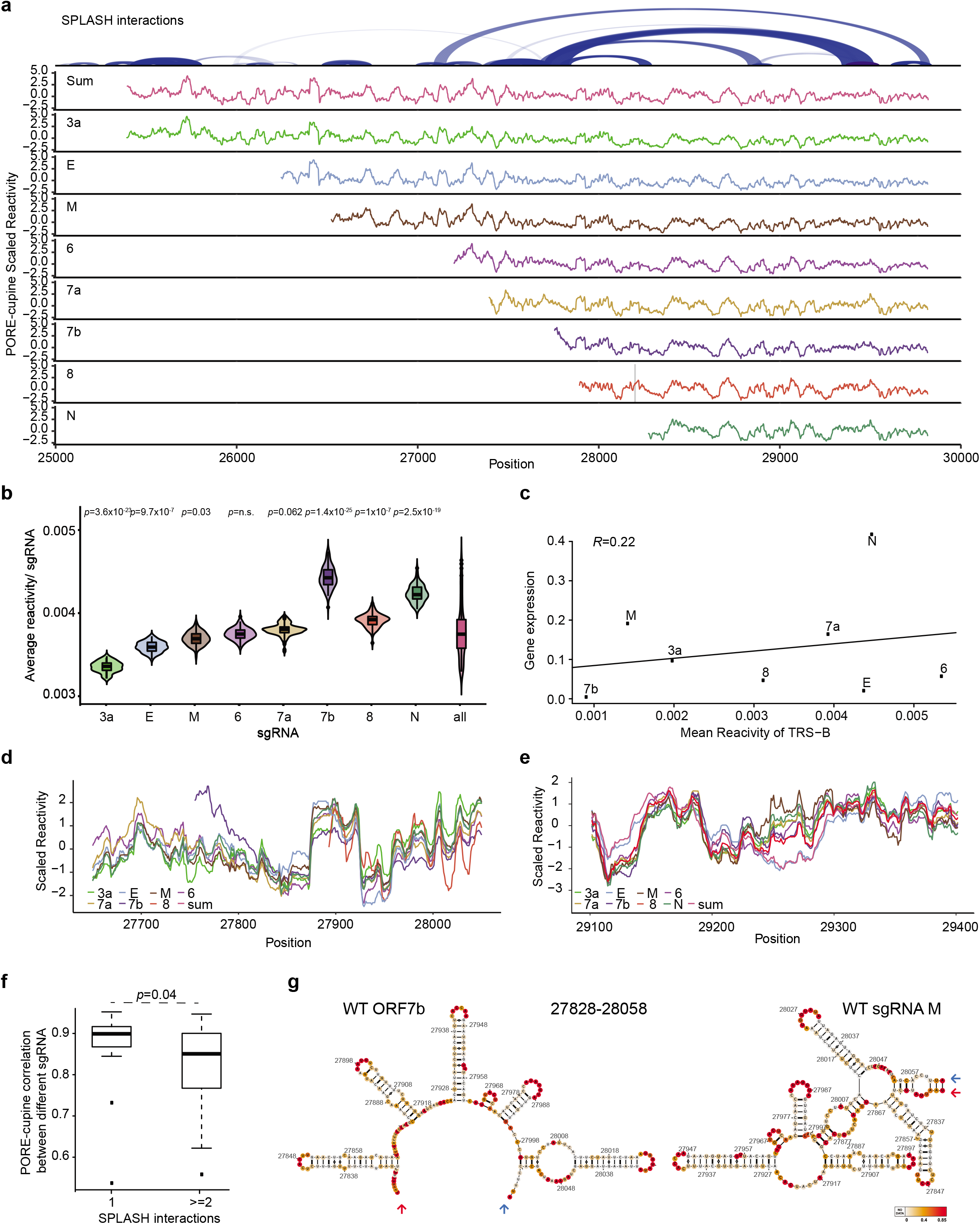
PORE-cupine reveals subgenomic RNA-specific structures. **a**, *Top*, SPLASH interactions along SARS-CoV-2 from the region 3a to the 3’UTR. *Bottom*, PORE-cupine reactivity signals are averaged across all the signals from the subgenomic RNAs (Sum). PORE-cupine reactivity signals are also shown for 3a (green), E (blue), M (brown), 6 (purple), 7a (light brown), 7b (navy), 8 (red) and N (dark green). PORE-cupine reactivity signals for each sgRNA are filtered for full length sequences that contain leader sequences for each sgRNA. Reactivities for the leader sequence are not shown. Regions with significant differences are highlighted in grey, (**Methods). b**, Violin plots showing the distribution of average reactivities for each sgRNA. Each sgRNA is subsampled for 500 strands before calculating its mean, n=100. *P*-values are calculated by comparing the distribution of the reactivities in each sgRNA against all of the sgRNAs. **c**, Scatterplot showing the correlation between the PORE-cupine reactivity around TRS-B for each sgRNA (x-axis) against transcript levels inside cells (y-axis). **d, e**, Reactivity plots of regions that show significant structure differences between the sgRNAs. **f**, Boxplots showing the distribution of correlation between reactivities of different sgRNAs for regions that show unique SPLASH interactions (1) and regions that show alternative SPLASH interactions (>=2). Regions that show alternative SPLASH interactions take on different conformations and show lower reactivity correlations between sgRNAs. **g**, Structure models of WT ORF7b and sgRNA M are generated using the program RNA Structure, using PORE-cupine reactivities as constraints. PORE-cupine reactivities are mapped onto the secondary structure models. The red and blue arrows indicate the same positions (start for red and end for blue) in the structure models.

We identified 237 and 187 intramolecular interactions along the WT and Δ382 genomes respectively (**Figure 3a, Supp. Tables 3**,**4**). SPLASH pair-wise interaction patterns are largely consistent between WT and Δ382 genomes, indicating the robustness of our method (**Figure 3a**). 45.6% and 42.3% of the intramolecular interactions occur over distance longer than 1kb in the WT and Δ382 genomes respectively, indicating that the viral sequences are involved in extensive long-range interactions (**Figure 3b, Supp. Figure 4c,d**). Longer-range interactions (>1kb) tend to have lower number of reads than shorter-range interactions, indicating that they are formed more transiently inside cells (**Figure 3c**), consistent with previous literature that longer-range interactions tend to be disrupted^32^. As RNA structures could have an impact on regulation of virus gene expression, we examined whether RNA pairing could be associated with translation using publicly available SARS-CoV-2 ribosome profiling data. We observed that ribosome pause sites from cycloheximide experiments have more pair-wise interactions than non-pause sites (**Figure 3d**)^33^, suggesting that RNA structures could be associated with translational pauses and thus regulate the translation of SARS-CoV-2.

Interestingly, we observed that SARS-CoV-2 RNA exhibit more alternative interactions than DENV and ZIKV RNAs inside the cell, with 55.6% and 48.1% of the WT and Δ382 pair-wise interactions involving two or more partners (**Figure 3e**)^10^, suggesting that SARS-CoV-2 RNA takes on numerous conformations that are present simultaneously inside host cells. We observed that a location at the 3’ end of sgRNA N is particularly promiscuous and interacts with regions throughout ORF1a (**Figure 3f**). Structure modelling of SPLASH identified interactions using the program RNA-cofold revealed that energies calculated from the predicted pairings are coherent with the SPLASH interaction counts (**Figure 3g**)^34^, indicating that the relative abundance of SPLASH counts between different interactions could serve as a proxy for the relative prevalence of these interactions inside cells.

### SARS-CoV-2 sgRNAs are structurally different

In addition to the synthesis of the full-length genomes, a nested set of 3’ co-terminal sgRNAs are made in SARS-CoV-2 infected cells using discontinuous RNA synthesis^9^ (**Supp. Figure 5a**). These sgRNAs range from 2-8 kb long, contain a leader sequence and are produced at different amounts. While SHAPE-MaP provides information on single nucleotide SHAPE along the genome, short-read sequencing makes it difficult to map structure information unambiguously to individual sgRNAs. As such, it is unclear if individual sgRNA contains unique structures that are different from these in the full length RNA genome and in other types of sgRNAs that could possibly be important for sgRNA-specific functions and regulation of their replication and translation.

**Figure 5.**
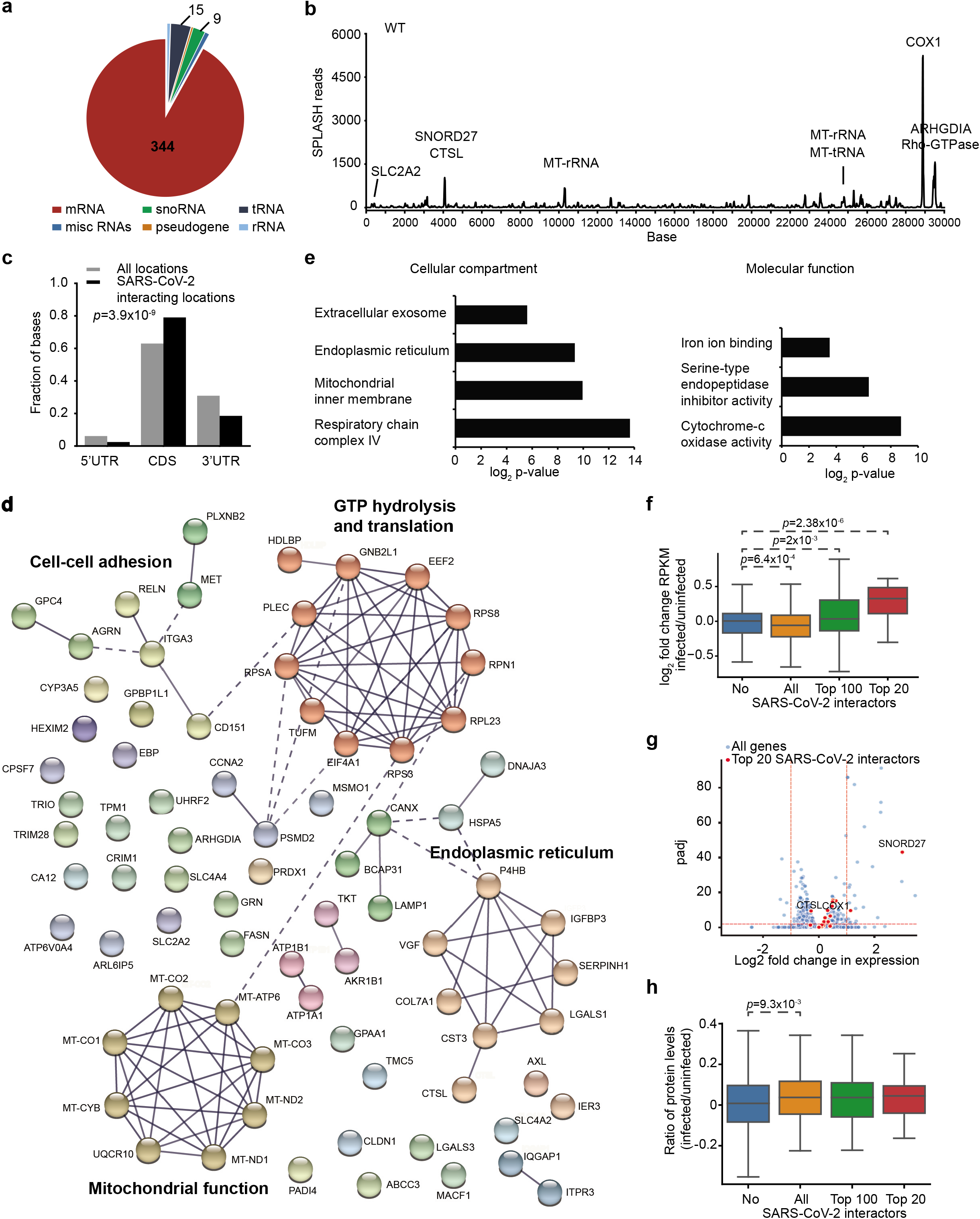
SARS-CoV-2 interacts with hundreds of host RNAs in vivo. **a**, Pie-chart showing the number of host RNAs from different RNA classes that interact with WT SARS-CoV-2. **b**, Line plot showing the number of SPLASH reads along the WT SARS-CoV-2 genome. The names of host RNAs that bind strongly to the virus at a particular location is labelled above the interaction peak. **c**, Bar-chart showing the fraction of host interacting regions that fall in 5’UTR, CDS and 3’UTR (black), as compared to what is expected from random (grey). Host interacting regions are enriched in CDS and depleted in 3’UTRs. **d**, STRING analysis of the top 25% SARS-CoV-2 host interactors^35^. The networks were built based on high confidence (0.7) evidence from protein-protein interaction sources of experiments, databases and text-mining where the line thickness indicates the strength of data support. Functional clusters in PPI networks were determined using Markov Clustering algorithm (MCL). The PPI enrichment *p*-value < 10^−16^. **e**, GO term enrichments of the top 25% SARS-CoV-2 interactors using David functional annotation analysis. SARS-CoV-2 interactors are enriched for transcripts that reside in the mitochondria, ER and exosome, and are enriched for molecular functions for iron binding, endopeptidase inhibitor activity and cytochrome-c oxidation activities. **f**, Boxplots showing the distribution of log2 fold change in gene expression upon SARS-CoV-2 infection in non-interacting genes, in 374 RNAs that interact with SARS-CoV-2 (All), in the top 100 interactors and top 20 interactors ranking by the chimeric read counts. SARS-CoV-2 interactors show a decrease in gene expression upon virus infection. However, the top interactors show an increase in gene expression upon virus infection, indicating that they are selectively stabilized. The expression data was calculated from the non-chimeric reads from SPLASH and quantified using DESeq2^57^. **g**, Volcano plot showing the distribution of host RNA gene expression upon SARS-CoV-2 infection. The top 20 interactors are highlighted in red and show a general stabilization in gene expression upon virus infection. **h**, Boxplots showing the distribution of protein ratio after virus infection in all genes, in 374 RNAs that interact with SARS-CoV-2, in the top 100 interactors and top 20 interactors.

To address this, we utilize our previously developed method of coupling RNA structure probing with Nanopore direct-RNA sequencing (PORE-cupine) to allow us to read out SHAPE reactivities along long RNA molecules^19^. Sequencing of two biological replicates of RNAs extracted from NAI-treated, WT and Δ382 SARS-CoV-2 infected Vero cells showed good structure correlation, indicating that our data is reliable (**Supp. Figure 5b**,**c**). We also confirmed that PORE-cupine reactivity shows good correlation with SHAPE-MaP reactivity along the SARS-CoV-2 genome (**Supp. Figure 5d**). By filtering for the full length reads that contain leader sequences, we determined reactivities along individual ORF3a, E, M, ORF6, ORF7a, ORF7b, ORF8 (WT only) and N transcripts (**Figure 4a, Supp. Figure 6a, Supp. Table 5**). We observed that ORF7b RNA contains the highest average reactivities for both WT and Δ382, suggesting that it is likely to be the most single-stranded among the different sgRNAs of SARS-CoV-2 (**Figure 4b, Supp. Figure 6b**). As structures around the leader sequences for each sgRNA were previously shown to have weak correlations with gene expression, we calculated the correlation between PORE-cupine reactivity around TRS-B sites for each sgRNA and their relative abundance from our Nanopore data. We observe a weak positive correlation between reactivity and transcript abundance, similar to previously published literature^28^, for both WT and Δ382 sgRNAs, suggesting that single-strandedness around the TRS-B region could result in increased synthesis of corresponding sgRNAs (**Figure 4c, Supp. Figure 6c**).

**Figure 6.**
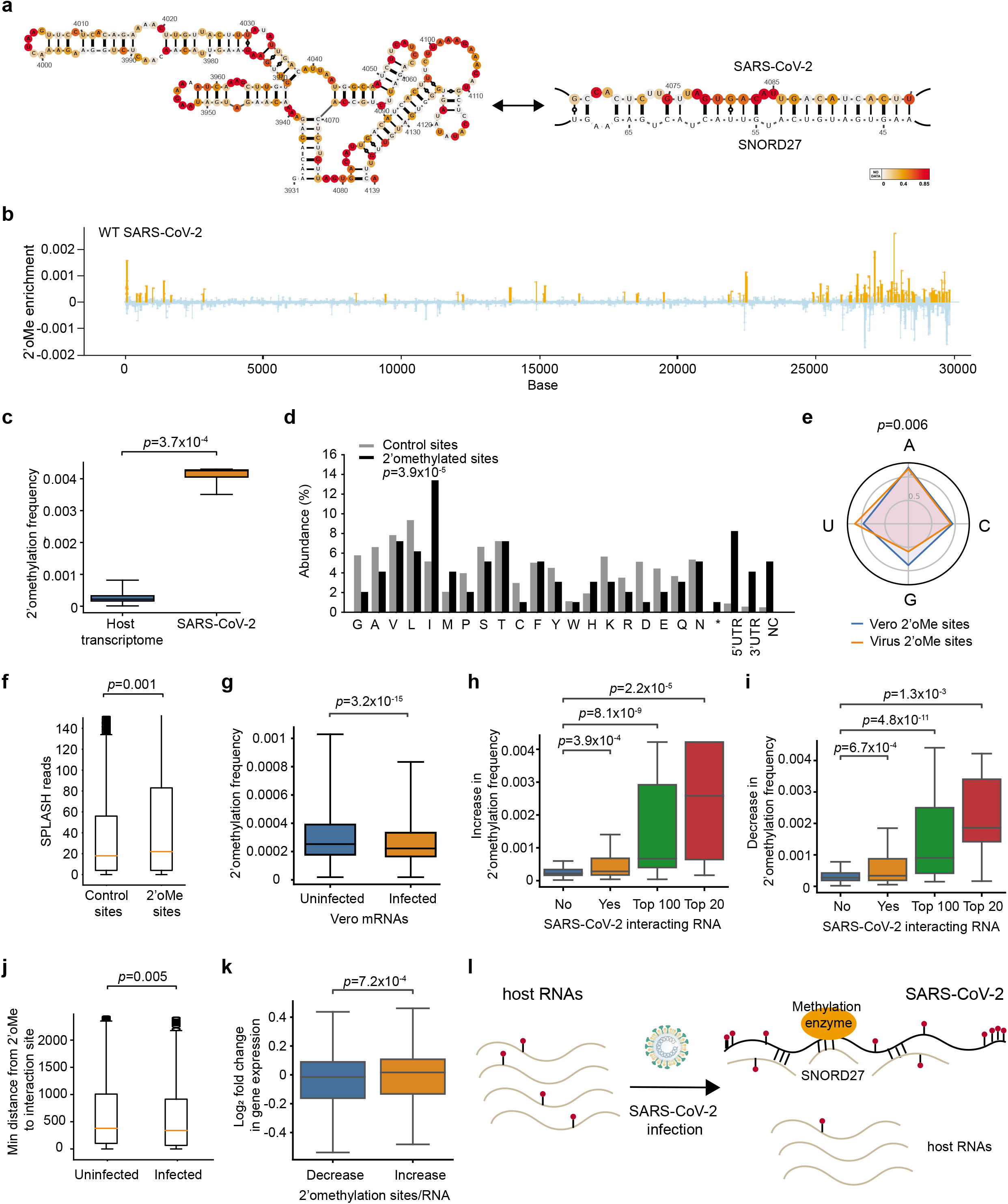
2’-O-methylation of SARS-CoV-2 sequesters methylation away from host RNAs. **a**, Structure model of SARS-CoV-2 before and after SNORD27 binding. The model of SARS-CoV-2 before SNORD27 binding is generated using the RNA structure program^53^ and incorporating SHAPE-MaP reactivity as constraints. The model of SARS-CoV-2:SNORD27 pairing is generated using RNAcofold^34^. SHAPE-MaP reactivities are mapped onto the structure models. **b**, The distribution of 130 2’-O-methylation sites along WT SARS-CoV-2 genome. The orange bars indicate enriched sites above control. **c**, Boxplots showing the distribution of 2’-O-methylation frequency along the host RNAs and on SARS-CoV-2 RNA from 4 replicates. **d**, Bar-plot showing the distribution of 2’-O-methylation sites (black), and control sites (grey), in the different amino acids, 5’UTR, 3’UTR and non-coding regions along the SARS-CoV-2 genome. *P*-value is calculated using chi-squared test. **e**, Distribution of 2’-O-methylation sites on SARS-CoV-2 genome (orange) and Vero transcriptome (blue) at A, C, U, G bases. The proportion of each nucleotide was normalized by its prevalence in the host transcriptome and SARS-CoV-2 genome respectively. 2’-O-methylation sites on SARS-CoV-2 are enriched in Us and depleted in Gs. *P*-value is calculated using chi-squared test. **f**, Boxplot showing the distribution of SPLASH chimeric reads at all sites versus 2’-O-methylation sites along the SARS-CoV-2 genome. *P*-value is calculated using Wilcoxon Rank Sum test. **g**, Boxplot showing the distribution of 2’-O-methylation frequency along Vero mRNAs in uninfected and SARS-CoV-2 infected cells. *P*-value is calculated using Wilcoxon Rank Sum test. **h, i**, Boxplot showing the distribution of increase (**h**) or decrease (**i**) in methylation frequency in non-interacting RNAs, in all interacting RNAs, in top 100 interacting RNAs and in top 20 interacting RNAs. *P*-value is calculated using Wilcoxon Rank Sum test. **j**, Boxplot showing the distribution of the minimum distance between host 2’-O-methylation site and the location of its interaction with SARS-CoV-2. *P*-value is calculated using Wilcoxon Rank Sum test. **k**, Boxplot showing the distribution of changes in gene expression in Vero mRNAs that either lost (decrease) or gained (increase) 2’-O-methylation sites. *P*-value is calculated using Wilcoxon Rank Sum test. **l**, Model of our hypothesis. SARS-CoV-2 binds to SNORD27 to sequester methylation enzymes to itself and away from host mRNA, enhancing host RNA decay.

To identify structures unique to each sgRNA, we compared the reactivities among individual sgRNAs to identify highly consistent as well as divergent structural regions (**Figure 4a, Methods**). We found 4 regions in RNAs of WT SARS-CoV-2 that showed consistent structure differences between different sgRNAs, 3 of which are also seen in the RNAs of Δ382 sgRNAs (**Supp. Figure 6a**). While two regions centred around bases 27800 and 28250 correspond to the leader sequences of sgRNAs of ORF7b and N respectively, two other structurally different regions (centred around 29300 and in 3’UTR) are present within all sgRNAs, and hence cannot be identified using short-read sequencing (**Figure 4d**,**e, Supp. Figure 6a**,**d**,**e**). We checked that the regions that show diverse structures in different sgRNAs also exhibit multiple interaction partners by SPLASH, confirming that those regions do exist in alternative conformations (**Figure 4f**). We then visualized the sgRNA-specific structures by incorporating PORE-cupine reactivities into structure modelling and observed different structure models for the same sequence region in different sgRNAs (**Figure 4g, Supp. Figure 7a**,**b**), further confirming that different sgRNAs could exist in different structures despite sharing the same sequences.

### Genomes of WT and Δ382 SARS-CoV-2 contain different RNA structures

Viruses that contain genomes with various ORF8 deletions have been found in patients around the world^7^, however the mechanisms behind how such deletions impacts the virus are still largely unknown. To determine whether virus phenotypes could be associated with structural differences, we performed correlations of SHAPE-MaP reactivities between the two genomes. As expected, structures in WT and Δ382 genomes are generally highly correlated (*R*=0.62, **Supp. Figure 8a**), although we do observe local structure differences at the deletion region of around base 28000 (**Supp. Figure 8b**,**c**). SPLASH analysis around the deletion region also revealed differences in pair-wise interactions between WT and Δ382, confirming the local structure rearrangements between the two viruses (**Supp. Figure 8d**).

As the deletion occurs around base 28000 (ORF8), it is present not only in the full-length genome but also in most of the sgRNAs (except for sgRNA of N, starts site of which is located downstream of the deletion). Due to the extensive amount of sequence similarity between the different sgRNAs, it is difficult to map uniquely to each individual sgRNA using short-read sequencing. Consequently, it remained unclear whether the structure differences between WT and Δ382 are present in all the sgRNAs or only between some specific sgRNAs. To determine which sgRNA shows reactivity differences between WT and Δ382 genomes, we compared the PORE-cupine reactivity profiles of individual WT and Δ382 sgRNA to each other (**Supp. Figure 9a**). While we could only detect very local reactivity differences immediately before and after the deletion site when all of the WT and Δ382 sgRNA reads are summed and compared to each other as an aggregate (similar reactivity profiles obtained using short-read Illumina sequencing), we observed additional structure differences when individual WT and Δ382 sgRNAs are compared to each other (**Supp. Figure 9a**,**b**,**c**). We observed the largest structure differences between WT and Δ382 in ORF3a and E sgRNAs (**Supp. Figure 9a**). We also consistently observed a second structurally different region between WT and Δ382 sgRNAs at the bases 29200-29400 (**Supp. Figure 9b**,**d**), indicating that the deletion could impact distal structures that are located more than 1kb away. As expected, we did not observe reactivity differences between N-gene sgRNAs of WT and Δ382 viruses as this sgRNA is transcribed using TRS located downstream of the deletion region. This finding indicates that the reactivity differences between other sgRNAs of WT and Δ382 viruses are likely to occur in cis due to the deletion and not due to factors that may act in trans (**Supp. Figure 9a**,**b**). As sgRNA of N is by far the most abundant sgRNA of SARS-CoV-2 and it did not show structure differences between WT and Δ382 viruses^9^, differences in the reactivity between the 29300 region in WT and Δ382 genomes were masked when an aggregate reactivity of all sgRNAs is used for comparison (**Supp. Figure 9a**,**b**). As such, using long-read sequencing to map RNA structures across sgRNAs can yield novel insights into sgRNA-specific RNA structures.

### SARS-CoV-2 genome interacts strongly with mitochondrial RNAs and snoRNAs

The genomes of RNA viruses can interact directly with host RNAs to facilitate or restrict viral infection. By analysing the SPLASH interactions between SARS-CoV-2 and host cell RNAs, we identified 374 and 334 host RNAs that interact with the WT and Δ382 SARS-CoV-2 genomes respectively (**Figure 5a**,**b, Supp. Figure 10a**,**b, Supp. Tables 6**,**7**). The host RNA-virus genome interactions are preferentially enriched in the coding regions along host mRNAs (**Figure 5c**). STRING analysis of the top 25% of SARS-CoV-2 interactors showed that they are enriched for proteins that physically interact with each other (PPI: *p*<10^−16^)^35^, including genes that are involved in the mitochondria, ER, GTP hydrolysis and translation processes (**Figure 5d**). GO term enrichment of interacting RNAs showed similar enrichments, confirming the importance of SARS-CoV-2 interactors in mitochondrial and ER function (**Figure 5e**)^36,37^.

While SARS-CoV-2 RNAs bind to more than 300 RNAs inside cells, we observed that the top 10 (2.6%) of the strongest interactors contributed to 17.5% and 24.1% of all WT and Δ382 binding events, indicating that the virus binds to them particularly strongly (**Figure 5b**). These strong interactors include mitochondrial RNAs such as the mRNA of COX1, a mitochondrially encoded cytochrome-c oxidase, mitochondrial rRNA, mRNA of ARHGDIA, a Rho-GTPase signaling protein and SNORD27, a snoRNA responsible for 18S ribosomal RNA methylation (**Supp. Figure 10c**). Previous study using RNA-GPS had shown that SARS-CoV-2 is localized to the mitochondria and the nucleolus^38^. SARS-CoV-2 infection also results in mitochondrial dysregulation^23,39^. Further experiments are needed to test whether the direct pairing between SARS-CoV-2 and mitochondrial RNAs contributes to mitochondrial dysregulation.

SARS-CoV-2 infection has been found to have an impact on almost every aspect of the host transcriptome to control virus and host gene regulation^40^. We observed a general decrease of RNA abundance of SARS-CoV-2 interactors upon virus infection. Interestingly, however, an opposite trend was observed for the strong interactors that were selectively stabilized and their abundance increased (**Figure 5f**,**g, Supp. Figure 10d**,**e**). qRT-PCR analysis of key interactors such as COX1 mRNA and MT-rRNA showed that these RNAs are indeed stabilized upon virus infection, confirming our RNA sequencing results (**Supp. Figure 10f-i**). Mining of published SARS-CoV-2 proteomics data revealed that proteins encoded by SARS-CoV-2 interactors were also preferentially translated and/or stabilized at the protein level as compared to proteins produced by non-SARS-CoV-2 interactors (**Figure 5h**)^41^. Thus, interaction with SARS-CoV-2 RNAs may confer a stabilizing effect on their overall gene regulation.

### SARS-CoV-2 RNA binds to SNORD27 and is 2’-O-methylated

SNORD27 is one of the strongest host interaction partners for SARS-CoV-2 RNA (**Figure 6a**) and is traditionally known to guide 2’-O-methylation of 18S ribosomal RNA^42^. As snoRNAs can bind and methylate cellular RNAs^43^, and methylation enzymes including fibrillarin (FBL), rRNA methyltransferase 2 and 3 (MRM2 and MRM3) have been found to be physically associated with SARS-CoV-2 genome^23^, we tested whether SARS-CoV-2 RNA could be 2’-O-methylated and whether host RNAs’ methylation levels are changed upon virus infection. We performed 2 biological replicates of Nm-seq on total RNA from Hela cells, as well as from uninfected and SARS-CoV-2 infected Vero-E6 cells (**Supp. Figure 11a, Supp. Tables 8**,**9, Methods**)^44^. Biological replicates of Nm-seq from both cell types show that they are well correlated, suggesting that Nm-seq data is reproducible (**Supp. Figure 11b**,**c**). Nm-seq analysis on human 18S rRNA accurately identified 36 out of 42 known 2’-O-methylation sites and had a high AUC-ROC curve of 0.96, suggesting that we are able to detect existing 2’-O-methylation sites accurately and sensitively (**Supp. Figure 11d**,**e**).

Using Nm-seq, we identified a total of 130 2’-O-methylation sites in SARS-CoV-2 genome (**Figure 6b**), and 4,931 sites in 4,142 transcripts in the Vero transcriptome (**Supp. Figure 11f**,**g**). We observed that a 2’-O-methylated host mRNA contains approximately 1.1 modifications per transcript in the Vero transcriptome, similar to methylated RNAs in Hela cells^44^. The majority of these host modification sites (60%) were located in the coding regions and were enriched for codons encoding charged amino acids (**Supp. Figure 12a**), as previously described^44^. In comparison, SARS-CoV-2 genome is 19X more modified than host mRNAs after normalizing for transcript length (**Figure 6c**). The 2’-O-methylations are enriched in the 5’ and 3’UTRs of SARS-CoV-2 (**Figure 6d**), depleted in position 2 of codons (**Supp. Figure 12b**), and are enriched for U and depleted for G bases along the genome (**Figure 6e**). 2’-O-methylation sites on SARS-CoV-2 are also associated with high SPLASH reads, indicating that they are located near positions with abundant intramolecular pair-wise interactions (**Figure 6f**).

As the modification of SARS-CoV-2 genome might sequester corresponding RNA modification enzymes away from the host transcriptome, we calculated the changes in modification rates in the host RNAs in the presence and absence of SARS-CoV-2 infection. We observed a decrease in host RNA 2’-O-methylation frequency upon virus infection, supporting our hypothesis that they become less methylated (**Figure 6g**). In addition, we also observed that host RNAs that interact strongly with SARS-CoV-2 genome show greater 2’-O-methylation changes, and show large losses and gains in modification sites within the RNAs (**Figure 6h**,**i, Supp. Figure 12c**). We hypothesized that RNAs interacting with SARS-CoV-2 genome could be methylated near their interacting regions, presumably due to proximity to SNORD27, while methylation sites that are located far away could be lost. To determine whether 2’-O-methylation sites on SARS-CoV-2 genome interacting RNAs are closer to the locations of virus-RNA interactions regions, we calculated the closest distance between virus-host interaction to host 2’-O-methylation site. We observed that sites that had 2’-O-methylation were indeed closer to virus-host RNA interactions sites (**Figure 6j**), supporting the hypothesis that proximity to SARS-CoV-2 genome might allow interacting RNAs to be methylated together within a hub.

2’-O-methylation has been shown to stabilize RNA gene expression inside cells^43^. Therefore, we hypothesized that the loss of 2’-O-methylation on host RNA upon virus infection may facilitate host RNA decay. Indeed, we observed that the abundance of host RNAs that show a decrease in methylation sites was significantly decreased in infected cells. In contrast, the abundance of host RNAs that show an increase in methylation sites was increased (**Figure 6k**). Thus, together with the production of NSP-1 from SARS-CoV-2 to cleave cellular RNAs, the binding of SARS-CoV-2 RNAs to SNORD27 could serve as part of a multi-prong mechanism to degrade cellular RNAs and maximize virus replication (**Figure 6l**)^45^.

## Discussion

Studying the molecular basis of virus pathogenicity enables us to understand how this can be counteracted and how to inhibit and target the replicating virus. By probing the local and pair-wise RNA interactions of the SARS-CoV-2 genome and sgRNAs using high throughput structure probing technologies on both the Illumina and Nanopore sequencing platforms, we identified potentially functional structure elements within the genome and demonstrated that these RNA structures are associated with ribosome pausing. While the structures along SARS-CoV-2 have also been probed using other short-read high throughput strategies such as DMS-sequencing^29^, these strategies have limitations in their ability to decipher sgRNA-specific structures due to extensive sequence similarity between the different sgRNAs. Using long-read sequencing, we identified both sgRNA-specific structures, as well as structure differences between WT and Δ382 genomes, that could serve as a basis for understanding sgRNA-specific functions in future.

While existing literature has mostly focused on understanding SARS-CoV-2: host protein interactions, here we describe that the virus genomes bind directly to hundreds of host RNAs inside cells using SPLASH. In addition to a previous report that showed that SARS-CoV-2 binds to snRNAs that are involved in splicing^46^, we identified diverse, functionally related, host mRNA-virus interactions and found that SARS-CoV-2 binds particularly strongly to mitochondrial RNAs and snoRNAs. Our results are consistent with previous predictions of SARS-CoV-2 localization in the mitochondria and nucleolus^38^, and the observation that the mitochondria is dysregulated upon SARS-CoV-2 infection^23^. In addition, previous studies have also shown that SNORD27 and mitochondrial RNAs are enriched on SARS-CoV-2 using formaldehyde crosslinking, and sequencing^23^. However, it is unclear if these RNAs bind to the genome directly or indirectly through protein interactions. Our studies show that SARS-CoV-2 RNA pairs directly with mitochondrial RNAs and snoRNAs at specific locations on host RNAs and SARS-CoV-2 genomes. As snoRNAs recruit 2’-O-methylation modifications on their target RNAs, we additionally observed that SARS-CoV-2 genome is extensively 2’-O-methylated inside cells. This deepens our functional understanding of SARS-CoV-2 biology from the previous observation that SARS-CoV-2 interacts with methylation enzymes, including FBL, inside cells.

2’-O-methylation of RNA plays important roles inside cells and can contribute to RNA stabilization as well as key functions in innate immunity^47^. A recent study revealed that HIV-1 hijacks cellular proteins to 2’-O-methylate its genome to escape from host innate immune sensing by modulating the expression of type-1 interferons^48^. Further experiments are needed to determine if 2’-O-methylation of SARS-CoV-2 RNA could also allow it to escape host immunity. Another hypothesis for the binding of SARS-CoV-2 to SNORD27 is that the virus sequesters SNORD27 and methylation complexes towards itself and away from the rest of the Vero cell transcriptome. We observed that 2’-O-methylation sites on SARS-CoV-2 are enriched for paired interactions, in agreement with previous literature that 2’-O-methylation in HIV stabilizes alternative pairing conformation of the transactivation response element^49^. Importantly, we observed that 2’-O-methylation levels decrease in host RNAs after SARS-CoV-2 infection, supporting our hypothesis that the binding of SNORD27 to SARS-CoV-2 could direct methylation enzymes to SARS-CoV-2, and away from host RNAs. In addition to the function of SARS-CoV-2 NSP1 to degrade host RNAs^45^, this could serve as part of a multi-prong strategy for the virus to degrade host RNA for its own survival. Further experiments would be needed to definitively prove this hypothesis.

In summary, our study identifies new potentially functional structures along the SARS-CoV-2 virus, new host factors and alternations of host gene regulation upon SARS-CoV-2 infection, providing critical new understanding of SARS-CoV-2 pathogenicity.

## Methods

### Cells and viruses

African green monkey kidney, clone E6 (Vero-E6) cells (ATCC# CRL-1586) were maintained in Dulbecco’s modified Eagle Medium (DMEM) supplemented with 5% fetal bovine serum (5% FBS). SARS-CoV-2 wild-type (WX56) and Δ382 mutant (CA001) were isolated from COVID-19 patients in Singapore, as reported previously^7^.

### SHAPE-MaP structure probing of SARS-CoV-2 virus in Vero E6 cells

Vero E6 cells were infected with SARS-CoV-2 viruses (WT and Δ382) at a multiplicity of infection (MOI) = 0.01 for 1 h at 37°C. Following 1 h infection, virus inoculum was removed and replaced with DMEM-5% FBS. Flasks were incubated for 48 h at 37°C, 5% CO_2_.

At 48 hpi, cells were washed once with PBS and trypsin was added to detach the cells from the flask. The cells were collected and centrifuged at 2000 rpm. The pellet was resuspended in PBS and the cells were then separated into three reactions: (1) added 1:20 volume of 1M NAI (03-310, Merck, 25 µl of NAI in 500 µl of infected cells) and incubated for 15 min at 37 °C for structure probing; (2) added 1:20 volume of dimethyl sulfoxide (DMSO) and incubated for 15 min at 37 °C, as negative control; and (3) set aside a third portion of the infected cells without any treatment, for the denaturing control in the downstream library preparation process. The total RNA was extracted from the cells using E.Z.N.A. Total RNA Kit (Omega bio-tek) according to the manufacturer’s instructions. We then performed library preparation following the SHAPE-MaP protocol to generate cDNA libraries compatible for Illumina sequencing^18^.

### Interactome mapping of SARS-CoV-2 virus in Vero E6 cells

Vero E6 cells were infected with SARS-CoV-2 viruses (WT and Δ382) at a multiplicity of infection (MOI) = 0.01 for 48 h. The cells were washed once with PBS and trypsin was added to detach the cells from the flask. The cells were collected and centrifuged at 2000 rpm. The pellet was resuspended in PBS and the cells were then incubated with 200 µM biotinylated psoralen and 0.01% digitonin in PBS for 10 min at 37 °C. The cells were spread onto a 10cm dish and irradiated at 365 nm of UV on ice for 20 min. The cells were collected, and the total RNA was then extracted using E.Z.N.A. Total RNA Kit (Omega bio-tek) according to the manufacturer’s instructions. We performed SPLASH libraries similarly to the published protocol^20,50^.

### Direct RNA sequencing using Nanopore

Unmodified and NAI-treated total RNA from WT and Δ382 SARS-CoV-2 infected Vero E6 cells were sequenced using Nanopore direct RNA sequencing 002 kit. The samples are sequenced and aligned according to the method used by Kim et al^9^. We used EPI_ISL_407987 and EPI_ISL_414378 as the reference for WT and Δ382 strain, respectively.

### Nm-Seq library construction

To map RNA nucleotides with 2’-O-methyl modification, Nm-Seq^51^ was applied to total RNA of Hela, Vero E6 and Vero E6 infected with SARS-CoV-2. In brief, eight rounds of oxidation-elimination-dephosphorylation (OED) were performed to iteratively eliminate non-modification nucleotides from the 3’ ends of fragmented RNAs, the 2’-O-methylated nucleotides resist the OED and make the 3’ end of reads enriched at the 2’-O-methylation sites. Two biological replicates of SARS-Cov-2 infected Vero E6 (total 4 replicates) and uninfected Vero E6 (total 2 replicates) were used to generate Nm-Seq libraries. Ten ug of each samples were used following NmSeq protocol and NEBNext Small RNA library kit with revised customized adaptors as previously described^44,51^. For input control library without OED, 1ug of RNA were used. Libraries were multiplexed and subjected to high-throughput sequencing using Illumina Next-Seq Hi.

### Analysis of SHAPE-MaP experiments

Sequencing reads obtained from two replicates of SHAPE experiments were aligned with the respective sequences for the strains (WT: EPI_ISL_407987, Δ382: EPI_ISL_414378) and SHAPE values for each position calculated using =Shapemapper-2.15’ of Weeks et al^52^. Read depths obtained in the sequencing experiments allowed for conclusive determination of SHAPE reactivites at approximately 80% of positions. A reference alignment of WT and Δ382 sequences was obtained using =mafft-7.453’ with the L-INS-I strategy. Subsequently, local correlation of SHAPE reactivity between replicates and between WT and Δ382 was calculated using Pearson correlation. As Pearson correlation between replicates was >0.9, replicates were pooled for subsequent analysis.

### Modeling of global RNA structure

Using the above SHAPE reactivities and reference sequences, separate global RNA structure models for WT and Δ382 using =Superfold’ with a maximum base pairing distance of 600 nt, and default SHAPE slope (1.8) and intercept (−0.6) parameters^53^. Particular regions of interest such as the frame-shift element were modelled separately in a local context to search for pseudoknot structures using the =RNAstructure’ tools =partition-smp’ and =ProbKnot-smp’.

### Modeling of individual subgenomic RNA structure from Nanopore data

The distributions of mutation rates obtained from Nanopore RNA sequencing were compared with the distribution of SHAPE reactivity values from short-read based experiments described above. We found that a scale factor of 100 brings the mutation rate distribution from the Nanopore experiment in line with distribution observed for conventional SHAPE experiments. Hence, we applied this scaling factor and then employed these as SHAPE data in the same =Superfold’ protocol as for the full-length models described above^53^.

### PORE-cupine analysis of direct RNA sequencing data

Filtering for full-length sub-genomic RNA: To separate full length aligned reads into their sub-genomic transcripts, we used two filtering conditions for all sgRNAs except for ORF6: 1, The aligned reads need to contain the leader sequence, 2. The aligned positions after the leader sequence must fall within ± 100 of the annotated sub-genomic sequences. For ORF6, we had to extend the second filter to −300 of the annotated sub-genomic sequence.

We calculated the reactivity for each subgenomic transcript by using PORE-cupine, with two adjustments to the analysis. 1. The length filter was removed, as only full-length transcripts were used for the analysis. 2. To reduce the amount of computing resources required, 20000 strands from each subgenomic transcripts in the unmodified libraries were randomly selected and used for the generation of models.

To determine the differences between reactivity, Wilcoxon Rank Sum test was applied to a 101 bases window with a step size of 25 nucleotides. Reactivity differences were compared across shared sequences between the different subgenomic transcripts within each strain (WT or Δ382), and across the two different strains. For the comparison between WT and Δ382 genome, the region in the WT strain that was not present in the Δ382 strain was masked. The *p*-values are corrected with Hommel’s method. In addition to using the statistical test to determine the differences, we added a second criteria of Pearson correlation < 0.7.

### Analysis of SPLASH experiments

Chimeric reads were divided into host-host, host-virus and virus-virus interactions for WT and Δ382 genomes. Virus-virus interactions were normalized to total virus-virus interactions and are shown in Figure 3A (WT: blue, Δ382: red). Virus-host interactions were equally normalized and the main locations for host interactions are shown in Figure 5B.

### Protein-protein interaction network analysis by STRING

The top 25% of host RNA interactors with SARS-CoV-2 were used as input for STRING analysis^35^, a search tool for retrieval of interacting genes to acquire protein-protein interaction (PPI) networks. The networks were built based on high confidence (0.7) evidence from protein-protein interaction sources of experiments, databases and text-mining where the line thickness indicates the strength of data support. Functional clusters in PPI networks were determined using Markov Clustering algorithm (MCL). The PPI enrichment *p*-value < 1.0e-16.

### Nm-Seq bioinformatics analysis

We referred to the pipeline from published protocol with some modifications^51^. The adaptors on raw reads were trimmed using Cutadapt and the reads without 3’ adaptors were discarded^54^. The PCR duplication reads were filtered out by the pentamers at 5’ and 3’ end of the reads as barcode using a custom script. The pentamers were then removed and reads that are shorter than 15 nt were discarded. The reads that passed all these filters were mapped to the reference which combined the longest transcriptome of *Chlorocebus sabaeus* (ensembl ChlSab1.1.101) and SARS-CoV-2 sequence (WT_EPI_ISL_407987s) by bwa-aln^55^. The multiple mapped reads and reads with soft and hard clipped alignments were discarded. The depth of each position on the transcriptome using 3’ end of reads were calculated in both input control and OED enriched libraries (D_input_, D_OED_). The depths were normalized by the read counts of each transcript. The significantly enriched sites over the other regions on the same transcript were detected by ΔD = D_OED_ − D_input_ using Z-test (FDR <0.05, D_OED_ >=10, D_OED_/D_input_ > 1.5).

As the SARS-CoV-2 transcripts comprise of 16∼30% of the total reads, we subsampled the two uninfected libraries into 4 libraries containing same reads on host transcriptome with the 4 infected replicates respectively, to compare the Nm sites of host transcriptome fairly. Bases that show 2’-O-methylation enrichment in 3 out of 4 replicates were recognized as the 2’-O-methylation sites.

## Acknowledgements

We thank members of the Wan lab for helpful discussions. YW is supported by funding from A*STAR, NRF, EMBO Young Investigatorship and CIFAR global fellow scholarship.

## Author contributions

Y. Wan conceived the project. Y. Wan, RG Huber, L.F. Wang, A. Lezhava and A. Merits designed the experiments and analysis. S.L. Yang, D.E Anderson, A. Aw, S.Y. Lim, X.N. Lim, K.Y.Tan, T. Chawla, Y. Su, P. Sessons performed the experiments. RG Huber, L.

DeFalco, Y. Zhang, A. Aw, T. Zhang performed the computational analysis. Y. Wan organized and wrote the paper with RG. Huber and all authors.

## Supplementary Figure Legends

**Supp. Figure 1. Quality matrixes for SHAPE-MaP for SARS-CoV-2. a**,**b**, Mutation rates (left) and read depth (middle) for modified, untreated and denatured WT (**a**) and Δ382 (**b**) SARS-CoV-2. Distribution of the SHAPE-MaP reactivities and standard error between replicates is shown on the right. **c**, Scatterplot of the SHAPE-MaP reactivity for 2 biological replicates of WT (left) and Δ382 (right). **d**, Violin plots showing the SHAPE-MaP reactivities in WT (left) and Δ382 SARS-CoV-2 for paired and unpaired bases.

**Supp. Figure 2. Structure modelling of SARS-CoV-2. a**,**b**, Structure models are generated using the program RNA structure, using SHAPE-MaP reactivities as constraints for the frameshifting element (**a**) and for the TRS-L elements along the genome (**b**). The SHAPE-reactivities are mapped onto the structure models. **c**, Table of base-pairing information in modelled RNA structures from WT, Δ382 SARS-CoV-2, DENV and ZIKV^10^.

**Supp. Figure 3. Highly structured and accessible regions along the SARS-CoV-2 genome. a**, Consensus regions (4/6 regions) along the SARS-CoV-2 identified by using different window sizes from 50 to 300 bases. **b**, Highly accessible (top 20% high SHAPE-reactivity regions) regions identified along WT and Δ382 SARS-CoV-2 genome (red). Regions that show high accessibility in both WT and Δ382 SARS-CoV-2 genomes are highlighted in purple. Leader regions and the positions of the different viral proteins are also shown along the genome.

**Supp. Figure 4. SPLASH identifies pair-wise RNA interactions within the SARS-CoV-2 genome. a**,**b**, 2D matrices showing the location of pair-wise intramolecular RNA interactions between 2 biological replicates of WT (**a**) and Δ382 (**b**) SARS-CoV-2 genomes. **c**, Structure models of pair-wise RNA interactions are generated using RNAcofold. SHAPE-MaP reactivities are mapped onto the modelled secondary structures. The SPLASH chimeric counts and predicted interaction energy for each model is also shown. **d**, Histogram showing the distribution of interactions that span different lengths along the Δ382 SARS-CoV-2 genome. Interactions longer than 1kb are classified as “long-range” and comprise 42.25% of all interactions.

**Supp. Figure 5. PORE-cupine reactivities provide information on subgenomic RNAs. a**, Schematic of the SARS-CoV-2 subgenomic RNAs expressed inside cells. **b, c**, Scatter plots of the PORE-cupine reactivities between 2 biological replicates for each subgenomic RNAs from WT (**b**) and Δ382 (**c**) SARS-CoV-2 genomes. d, Line plots showing the structure reactivity along SARS-CoV-2 obtained using SHAPE-MaP (red) and PORE-cupine (black). *R* (Pearson correlation) between the two plots is 0.49.

**Supp. Figure 6. PORE-cupine reactivities along the sgRNAs of Δ382 SARS-CoV-2. a**, PORE-cupine reactivity signals are averaged across all of the signals from the subgenomic RNAs (Sum). PORE-cupine reactivity signals are also shown for 3a (dark green), E (blue), M (brown), 6 (purple), 7a (light brown), 7b (navy), 8 (red) and N (dark green). PORE-cupine reactivity signals for each sgRNA are filtered for full length sequences that contain leader sequences for each sgRNA. Reactivies for the leader sequence are not shown. Regions with significant differences are highlighted in grey, **methods. b**, Violin plots showing the distribution of average reactivities for each sgRNA. Each sgRNA is subsampled for 500 strands before calculating its mean, n=100. *P*-values are calculated by comparing the distribution of the reactivities in each sgRNA against all of the sgRNAs.**c**, Scatterplot showing the correlation between the PORE-cupine reactivity around TRS-B for each sgRNA (x-axis) against transcript levels inside cells (y-axis). **d, e**, Line plots of the PORE-cupine reactivities of the different sgRNAs.

**Supp. Figure 7. Structure models of SARS-CoV-2 sgRNAs. a**,**b**, Structure models of WT ORF6 and ORF7a (**a**) and Δ382 sgRNA N and E (**b**) are generated using the program RNA Structure, using PORE-cupine reactivities as constraints^53,56^. PORE-cupine reactivities are mapped onto the secondary structure models. The red and blue arrows indicate the exact same start (red) and stop (blue) positions for the two structure models.

**Supp. Figure 8. SHAPE-MaP reactivity differences between WT and Δ382 genomes. a**, Scatterplot of SHAPE-MaP reactivities for WT and Δ382. *R* (Pearson correlation)= 0.621. **b**, Scatter plot showing read abundance (Y-axis) and Pearson correlation of SHAPE-MaP reactivity (X-axis). Regions with high read depth but low correlation are highlighted in red. **c**, Top, Line plot showing Pearson correlation of SHAPE-MaP reactivities between WT and Δ382 genomes. Bottom, read depth of the sequencing coverage along the genome. Regions that have high read depth and low correlation in terms of reactivities are shown in red. **d**, Arc plots showing the pair-wise RNA interactions around the Δ382 deletion site for WT (blue, top) and Δ382 (red, bottom) genomes.

**Supp. Figure 9. PORE-cupine reactivities differences between WT and Δ382 SARS-CoV-2 genomes. a**, PORE-cupine reactivity signals for WT (black) and Δ382 (red) are averaged across all the signals from their respective subgenomic RNAs (All reads). Line plots representing the sub-sampled lane are the averaged WT (black) and Δ382 (red) signals across the different sgRNA after the sgRNAs have been subsampled to the same depth, and hence carry equal weightage to each other. PORE-cupine reactivity signals for WT (black) and Δ382 (red) are also shown for sgRNAs 3a, E, M, 6, 7a, 7b, and N. PORE-cupine reactivity signals for each sgRNA is filtered for full length sequences that contain leader sequences for each sgRNA. Regions that show significant differences between WT and Δ382 sgRNAs are highlighted in blue, (**Methods**). **b**, Pearson correlation between WT and Δ382 PORE-cupine reactivities for sum of all sgRNAs for WT and Δ382 and between individual sgRNAs of WT and Δ382 genomes. **c**,**d**, Line plots showing the PORE-cupine reactivity of ORF6 along WT and Δ382 around the Δ382 deletion (between 27200-28600, **c**) and 1kb downstream of the deletion (29000-29600, **d**). **e**, Structure models of WT and Δ382 ORF6 are generated using the program RNA Structure, using PORE-cupine reactivities as constraints. PORE-cupine reactivities are mapped onto the secondary structure models.

**Supp. Figure 10. SARS-CoV-2: host RNA interactions. a**, Pie-chart showing the distribution of expressed RNAs in Vero cells (left) and RNAs that interact with Δ382 SARS-CoV-2 (right) in different RNA classes. **b**, Line plot showing the number of SPLASH reads along the Δ382 SARS-CoV-2 genome. The names of host RNAs that bind strongly to the virus at a particular location is labelled above the interaction peak. The pink box indicates the deletion region in Δ382. **c**, Structure models of pair-wise virus-host RNA interactions are identified using SPLASH and generated using the program RNAcofold. **d**, Boxplots showing the distribution of log2 fold change in gene expression upon Δ382 SARS-CoV-2 infection in all non-interacting genes, in 334 RNAs that interact with Δ382 (All), in the top 100 Δ382 interactors and top 20 Δ382 interactors. Δ382 SARS-CoV-2 interactors show a decrease in gene expression upon virus infection. However, the top interactors show an increase in gene expression upon virus infection, indicating that they are selectively stabilized. **e**, Volcano plot showing the distribution of host RNA gene expression upon Δ382 SARS-CoV-2 infection. The top 20 Δ382 interactors are highlighted in red and show a general stabilization in gene expression upon virus infection. **f-i**, qPCR results of host RNAs in Vero cells in the absence and presence of SARS-CoV-2.

**Supp. Figure 11. Nm-seq identifies 2’-O-methylation sites on SARS-CoV-2 and Vero transcriptome. a**, Schematic of the Nm-seq protocol to identify 2’-O-methylation sites in SARS-CoV-2 infected Vero cells. **b**, Scatter plot showing the distribution of 2’-O-methylation sites along 18S rRNA in 2 biological replicates of Hela total RNA. **c**, Scatter plot showing the distribution of 2’-O-methylation sites along SARS-CoV-2 in 2 biological replicates of virus infected Vero cells. **d**, The distribution of 36 identified 2’-O-methylation sites along human 18S rRNA. The orange bars indicate enriched sites above control. The red dots indicate known 2’-O-methylation sites. **e**, AUC/ROC curve of identified 2’-O-methylation sites on human 18S rRNA. **f**,**g** Venn diagrams showing the number of sites (**f**) and transcripts (**g**) that are 2’-O-methylated in Vero transcriptome with and without SARS-CoV-2 infection.

**Supp. Figure 12. Distribution of 2’-O-methylation sites on SARS-CoV-2 and Vero transcriptome. a**, Bar charts showing the distribution of 2’-O-methylation sites, and control sites, in different amino acids, 5’UTR, 3’UTR and non-coding regions on the Vero transcriptome. *P*-value is calculated using chi-squared test. 2’-O-methylation sites are enriched in charged amino acids. **b**, Bar charts showing the distribution of 2’-O-methylation sites along positions 1,2 and 3 of codons in the Vero transcriptome (left) and on SARS-CoV-2 (right). 2’-O-methylation sites are depleted on the 2^nd^ position of codons in SARS-CoV-2. *P*-value is calculated using chi-squared test. **c**,**d**, Distribution of 2’-O-methylation sites on COX1 (**c**) and CTSL (**d**) along its transcript, in uninfected (top) and infected (bottom) Vero cells.

