## Supplementary figures and images for "Comprehensive mapping of SARS-CoV-2 interactions in vivo reveals functional virus-host interactions"

### Supplementary Figure 1

##### Supplementary Figure 1

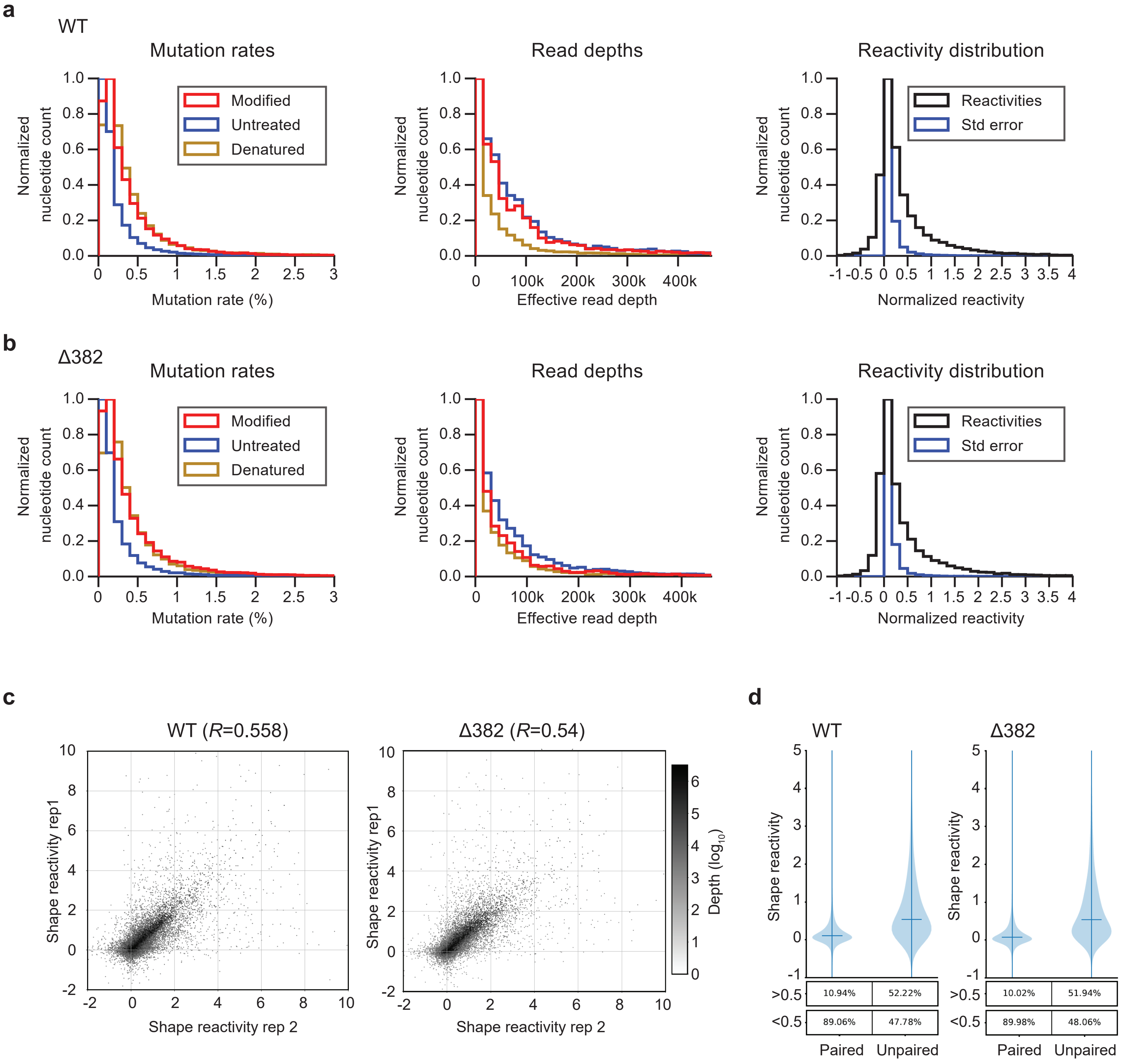

### Supplementary Figure 3

##### Supplementary Figure 3

**a**

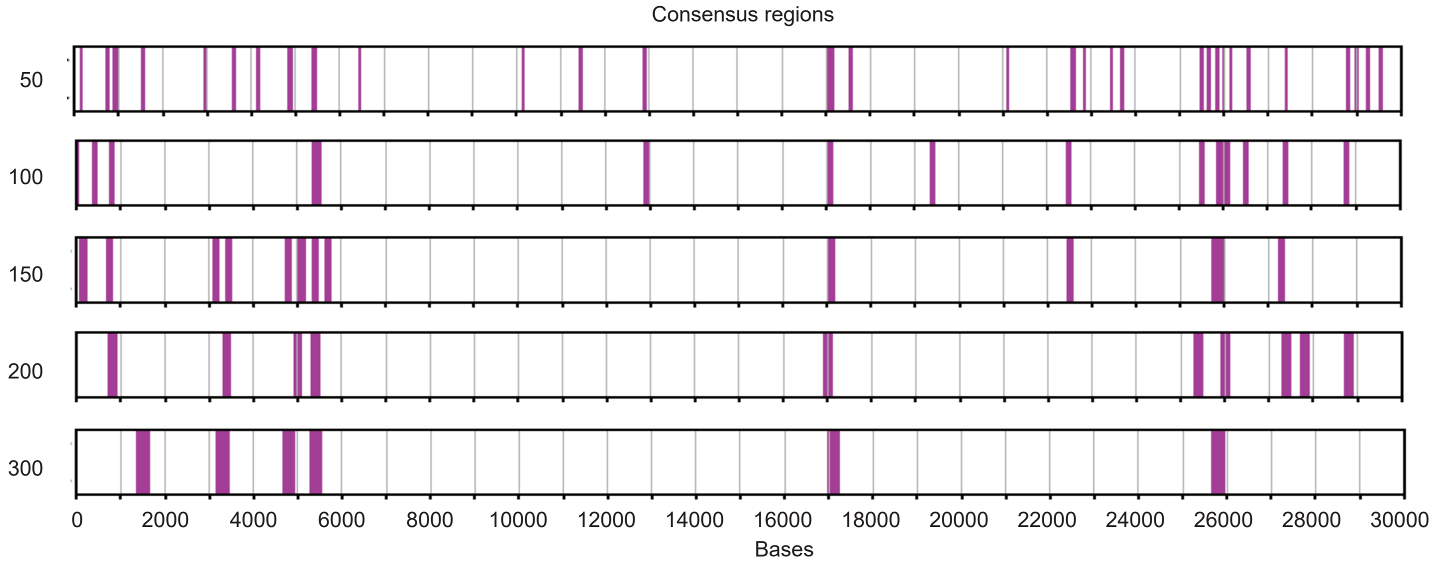


**b**

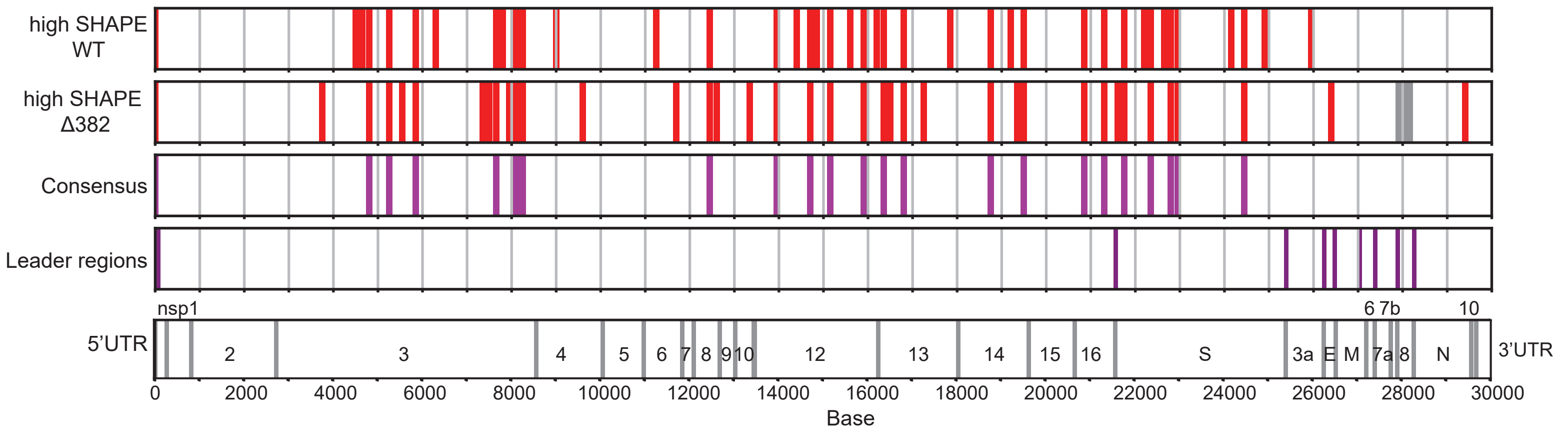

### Supplementary Figure 4

##### Supplementary Figure 4

**a**

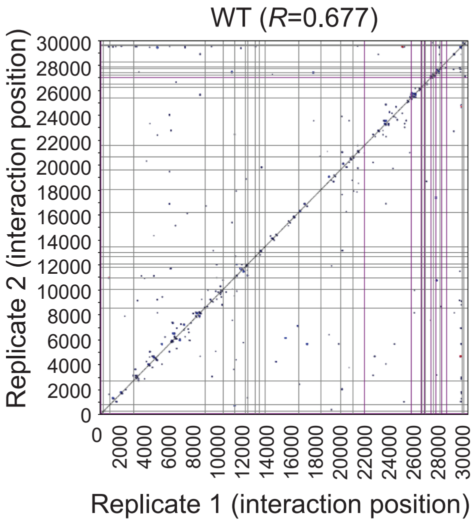


**b**

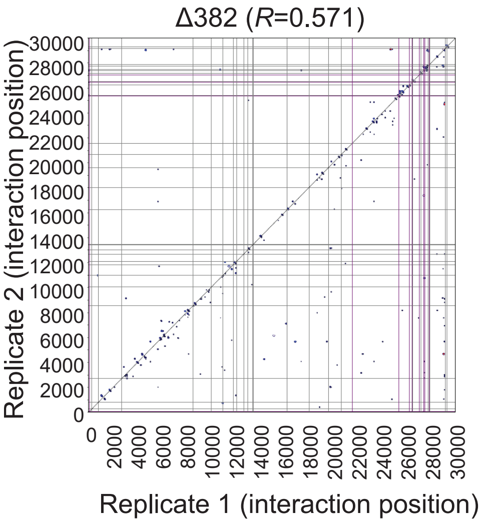


**c**

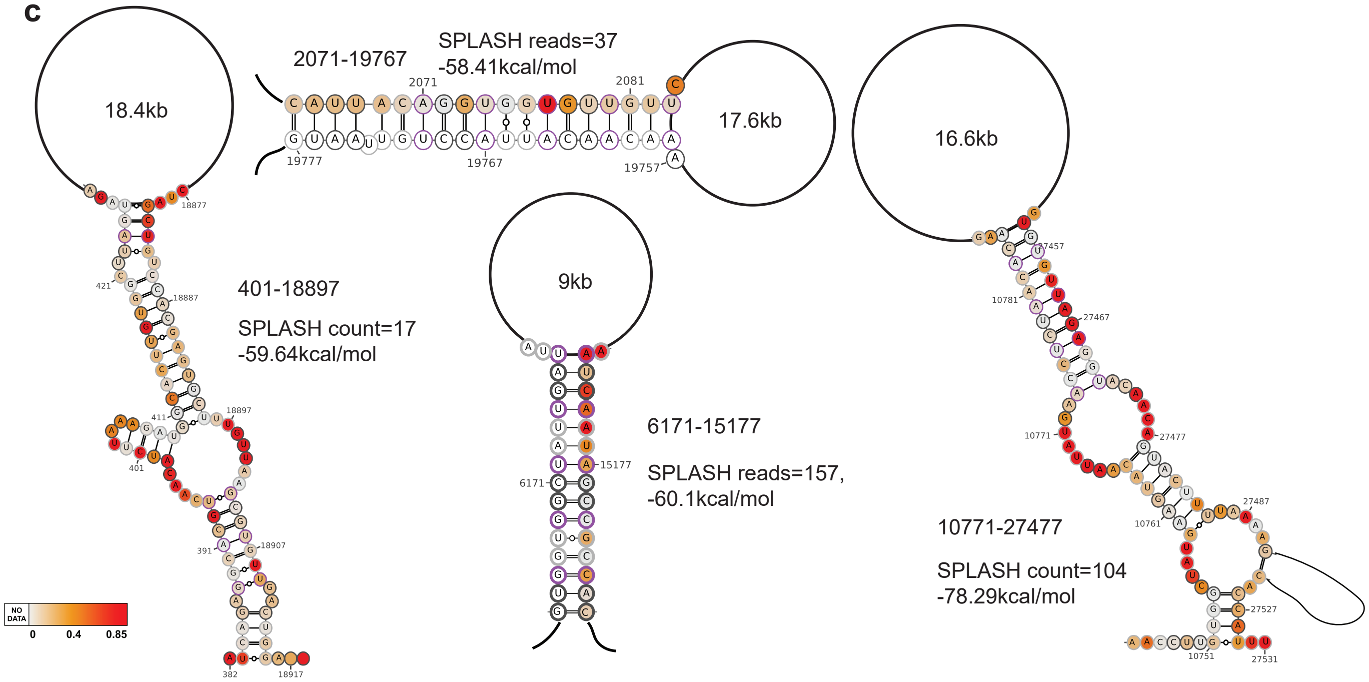


**d**

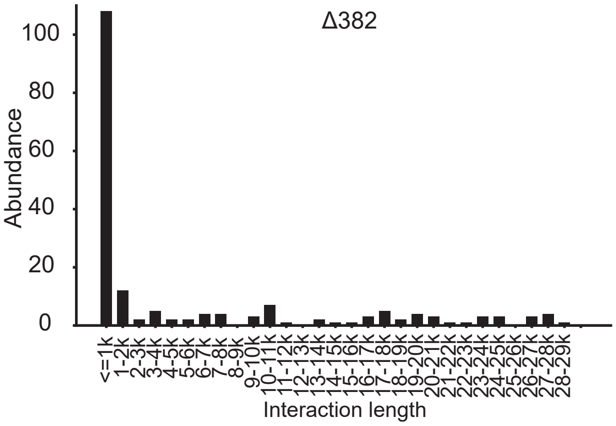

### Supplementary Figure 5

##### Supplementary Figure 5

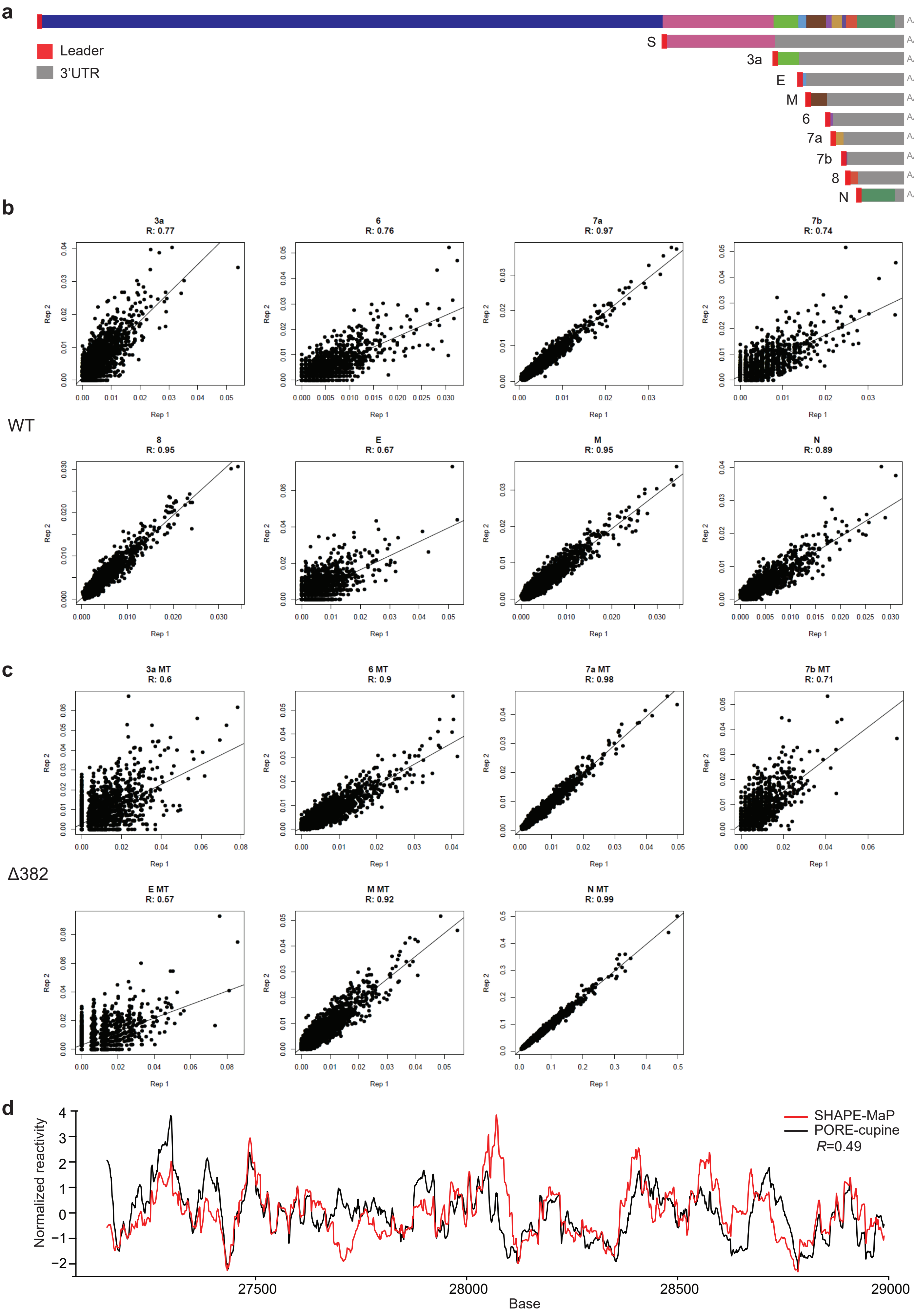

### Supplementary Figure 6

##### Supplementary Figure 6

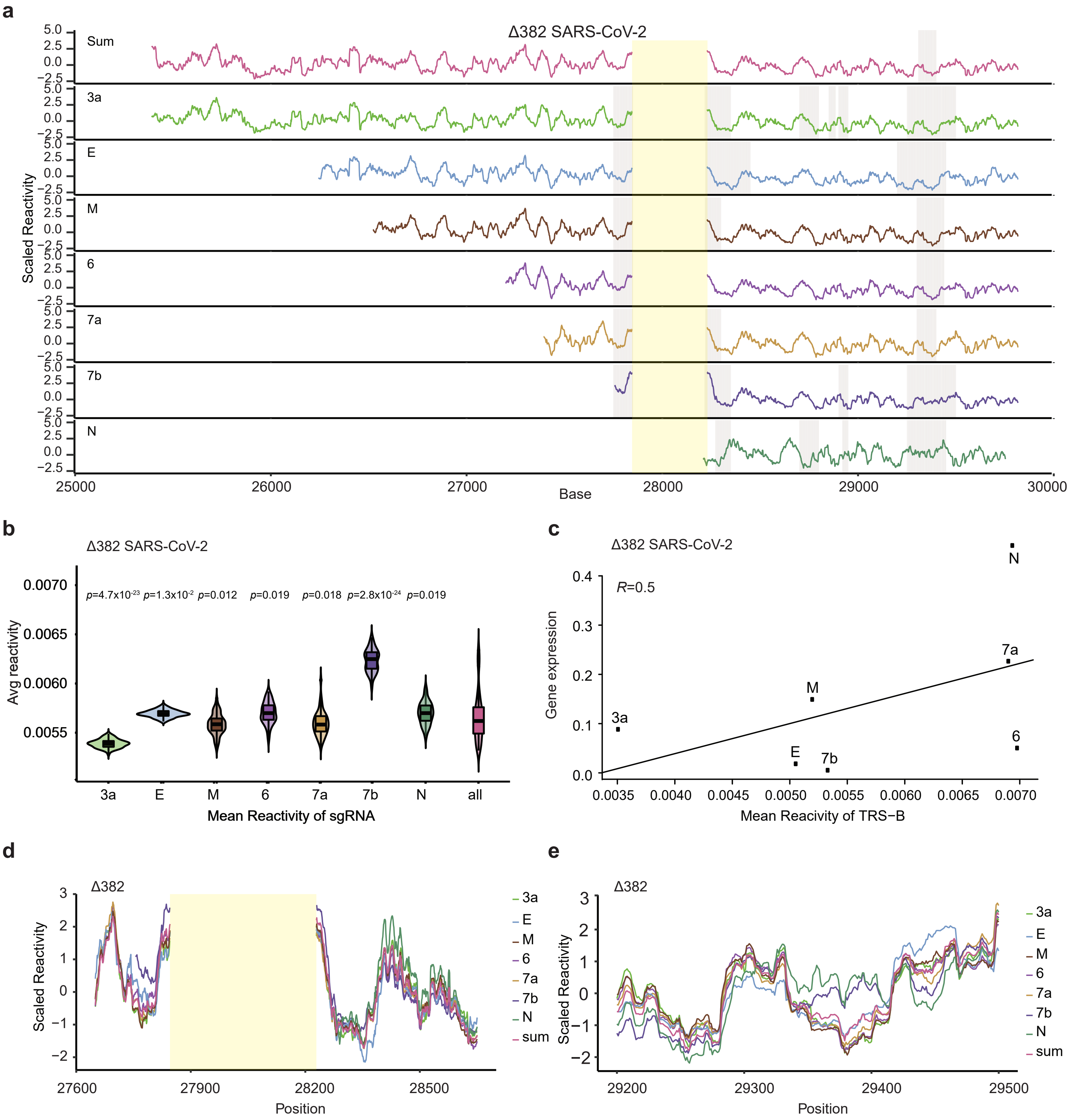

### Supplementary Figure 7

#### Supplementary Figure 7

**a**

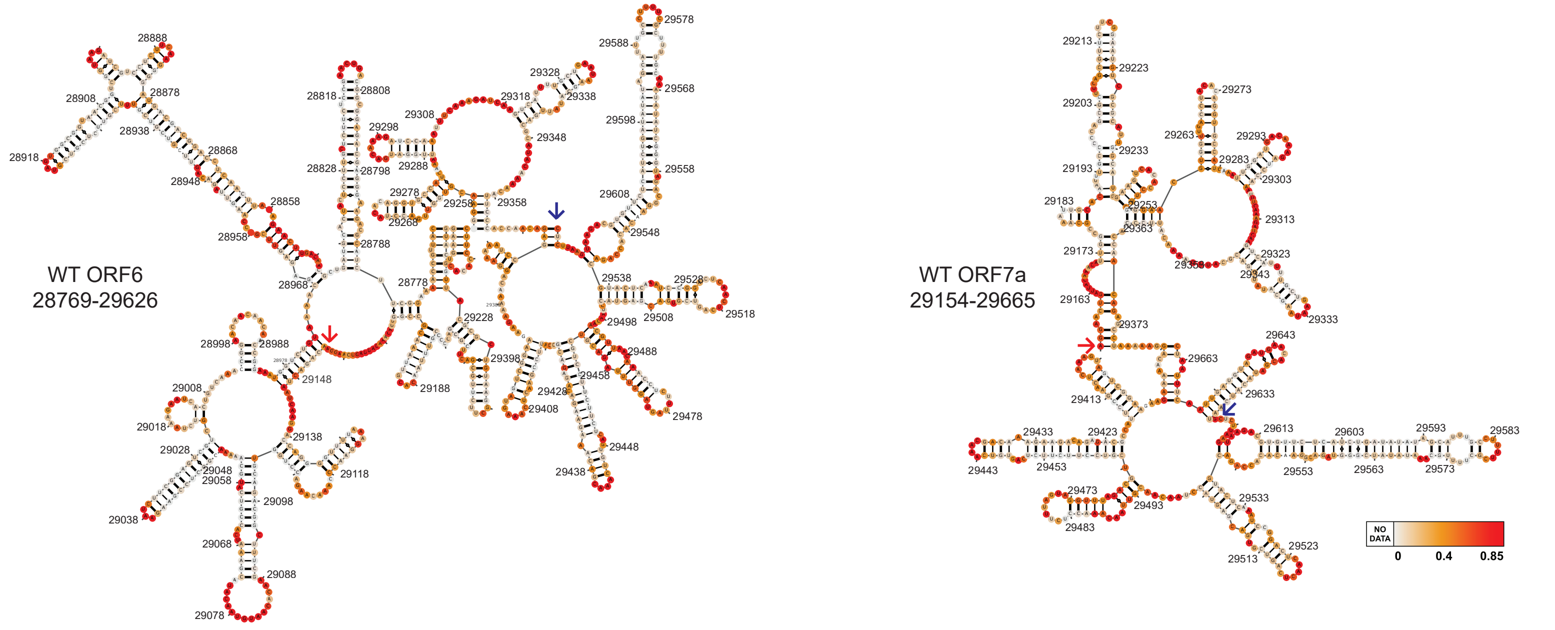


**b**

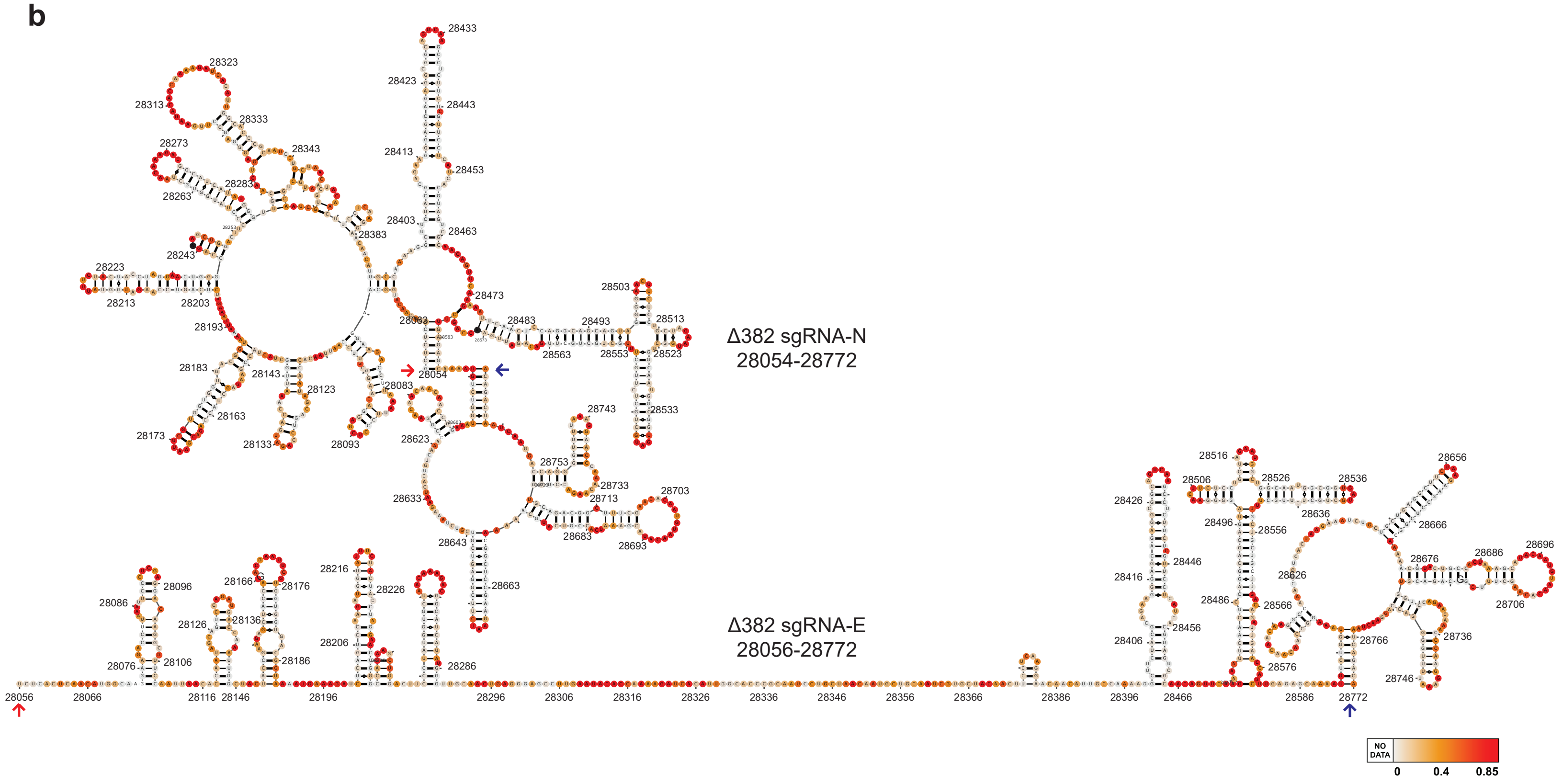

### Supplementary Figure 8

##### Suppmentary Figure 8

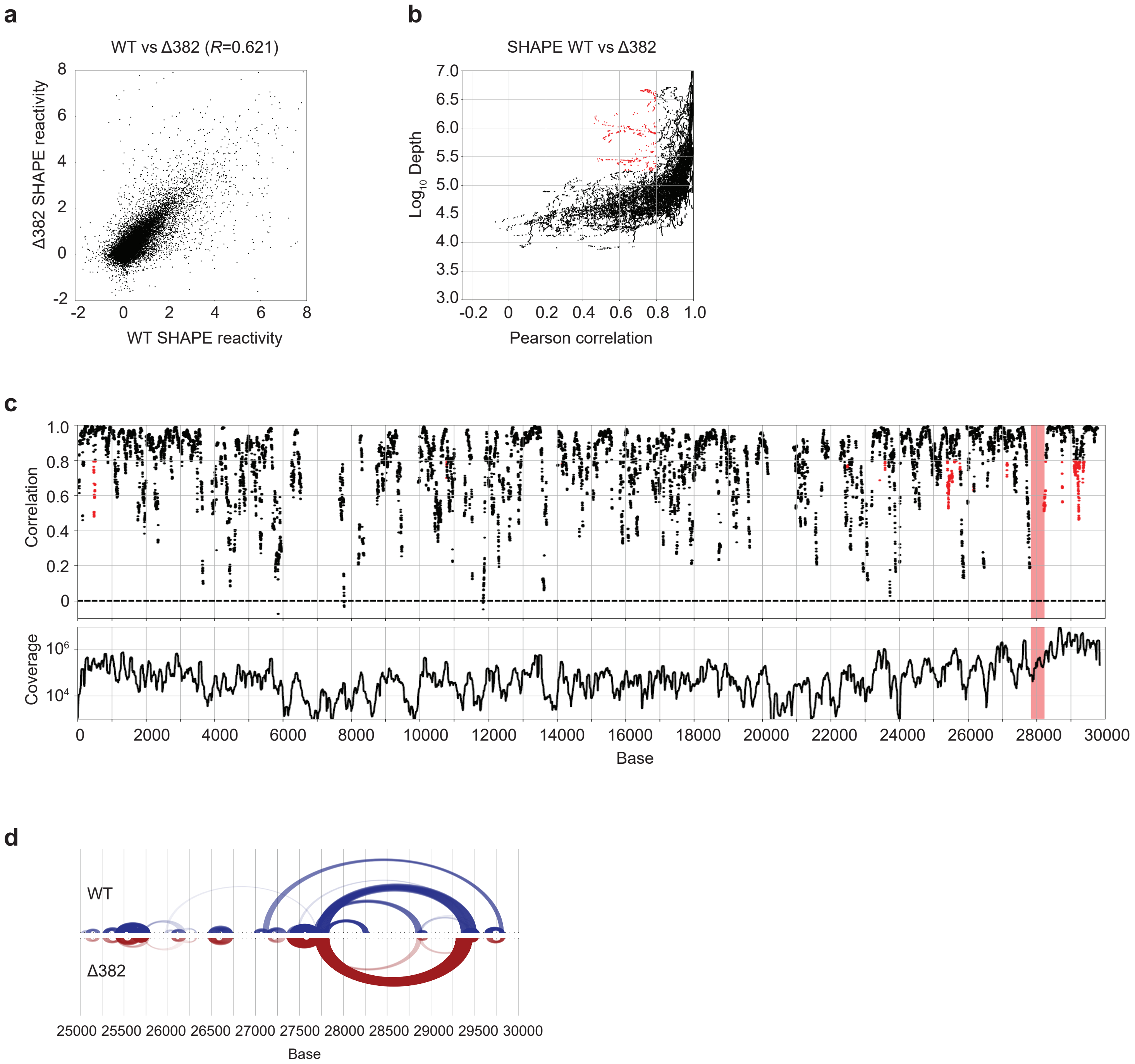

### Supplementary Figure 9

#### Supplementary Figure 9

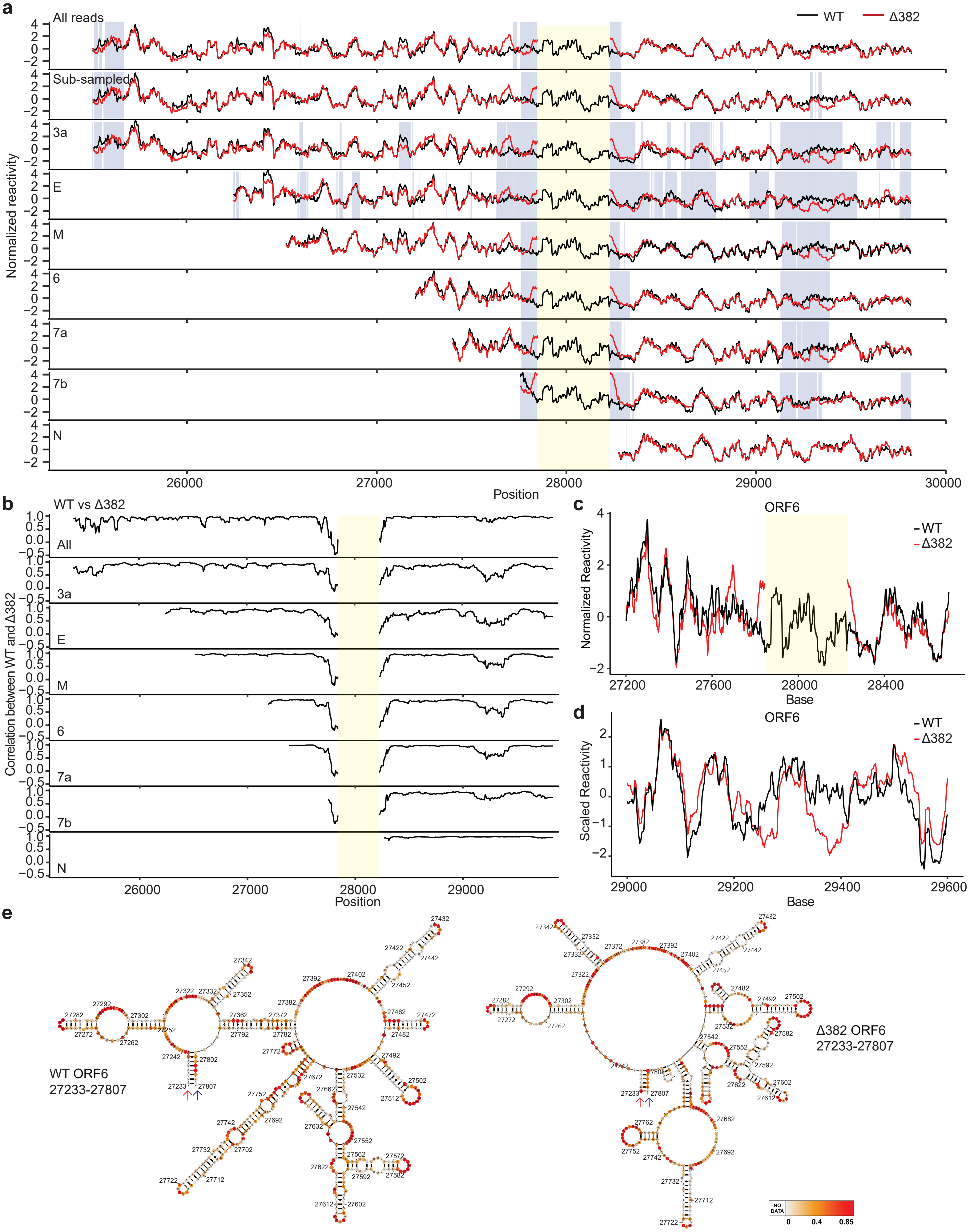

### Supplementary Figure 10

##### Supplementary Figure 10

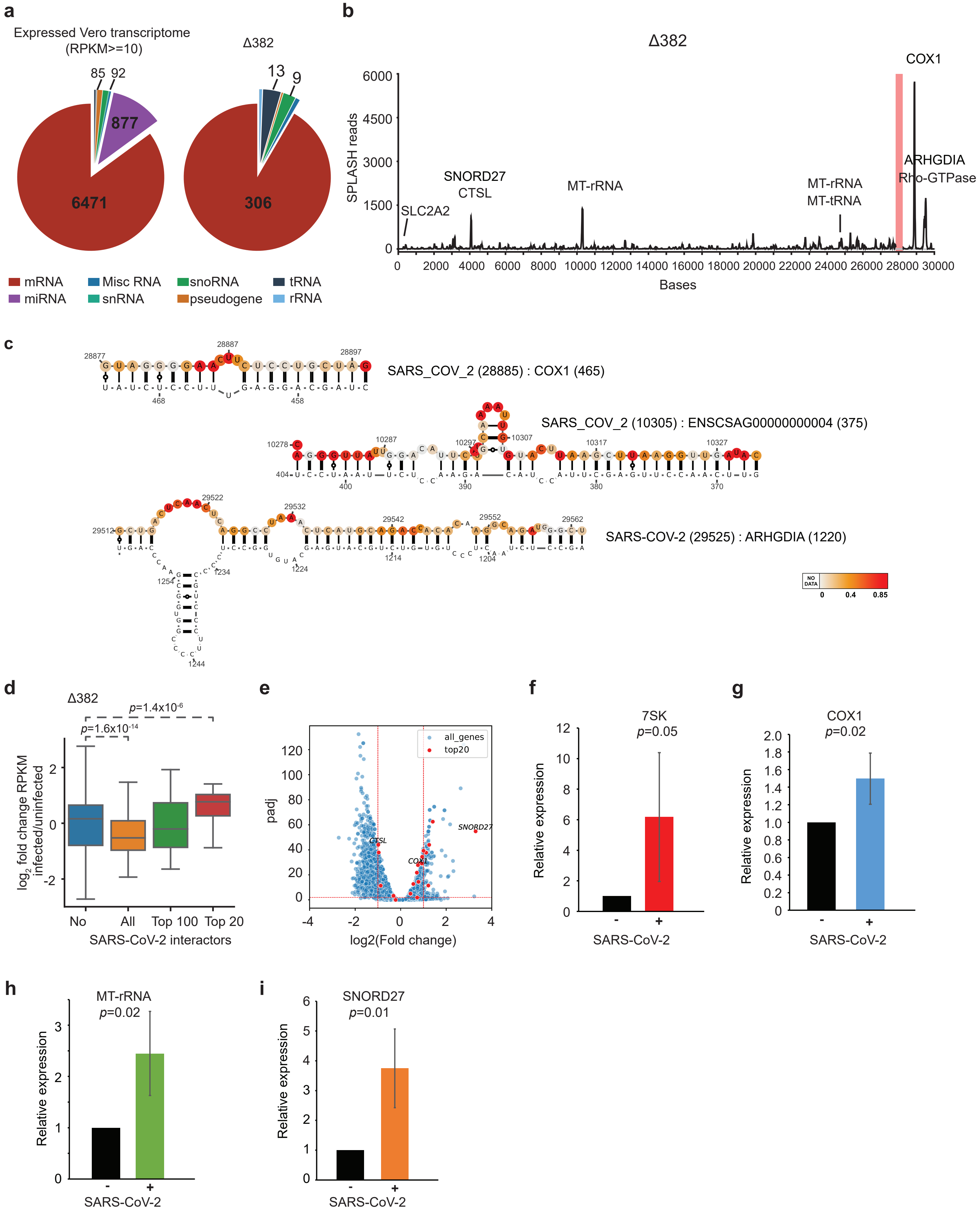

### Supplementary Figure 11

#### Supplementary Figure 11

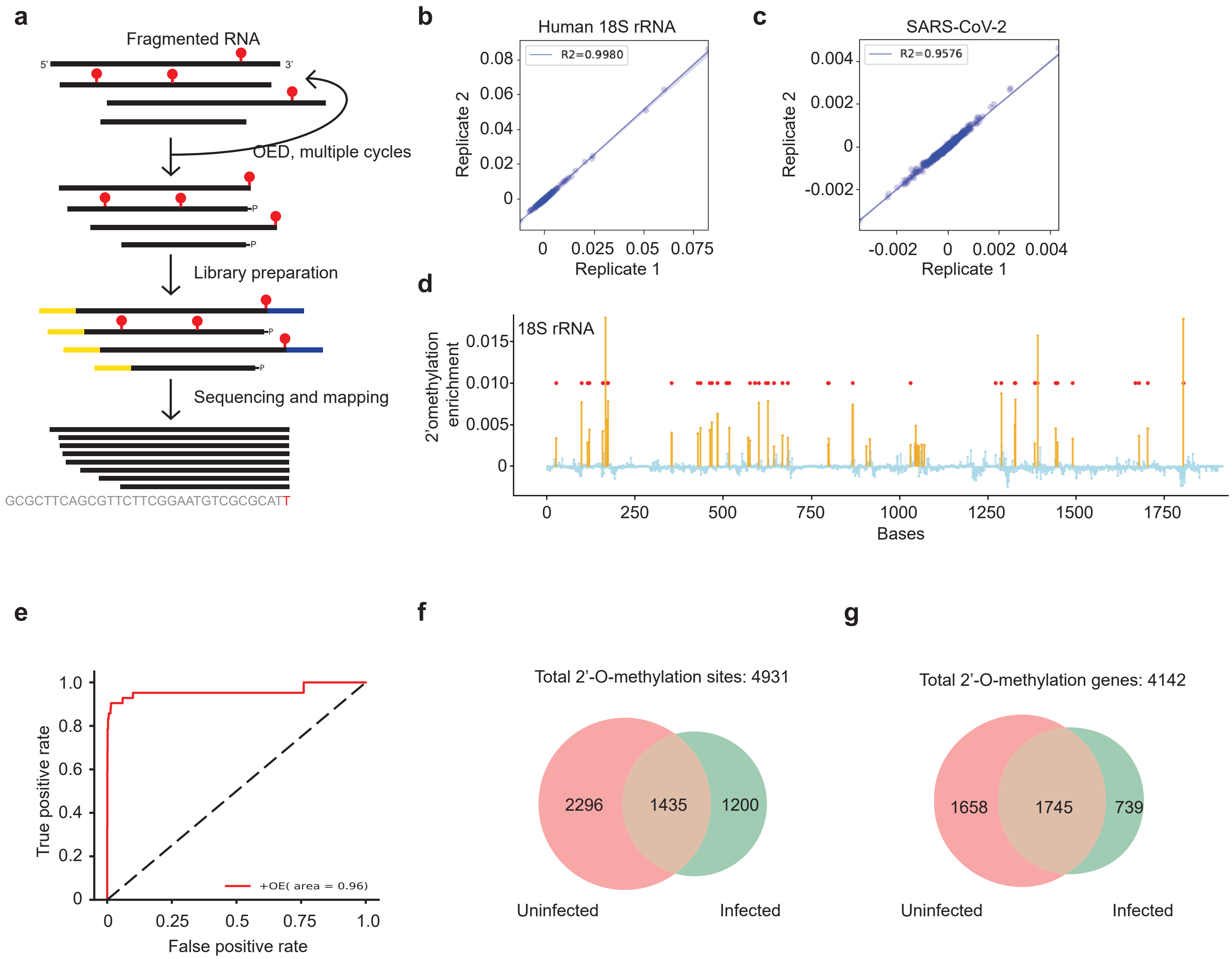

### Supplementary Figure 12

##### Supplementary Figure 12

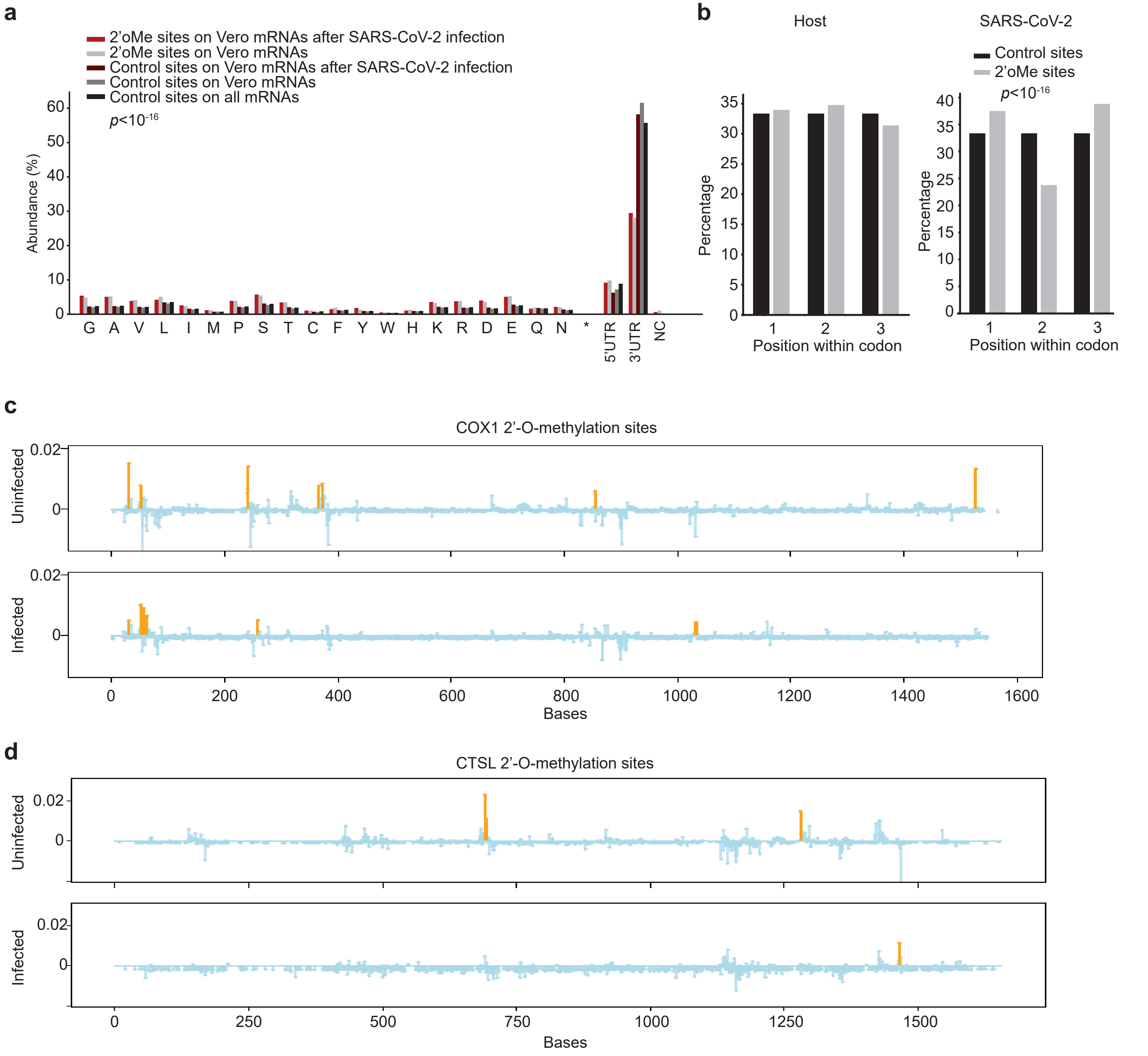
