## Supplementary Figure 2 for "Comprehensive mapping of SARS-CoV-2 interactions in vivo reveals functional virus-host interactions"

**a**

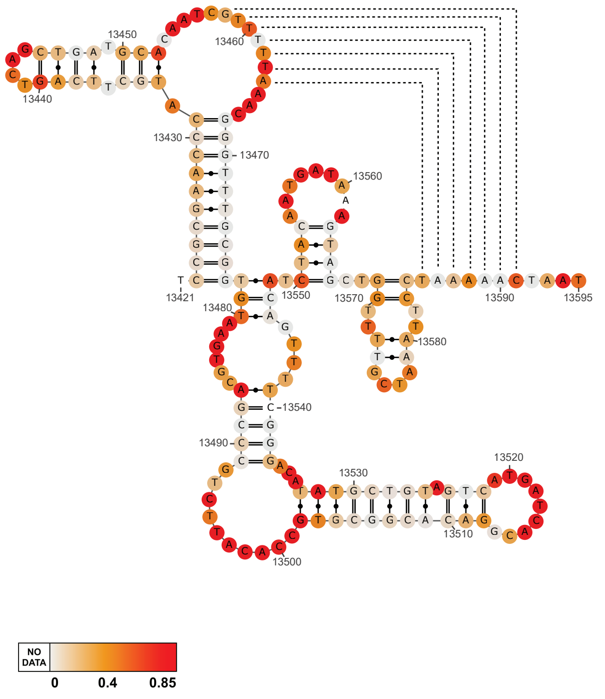


**b**

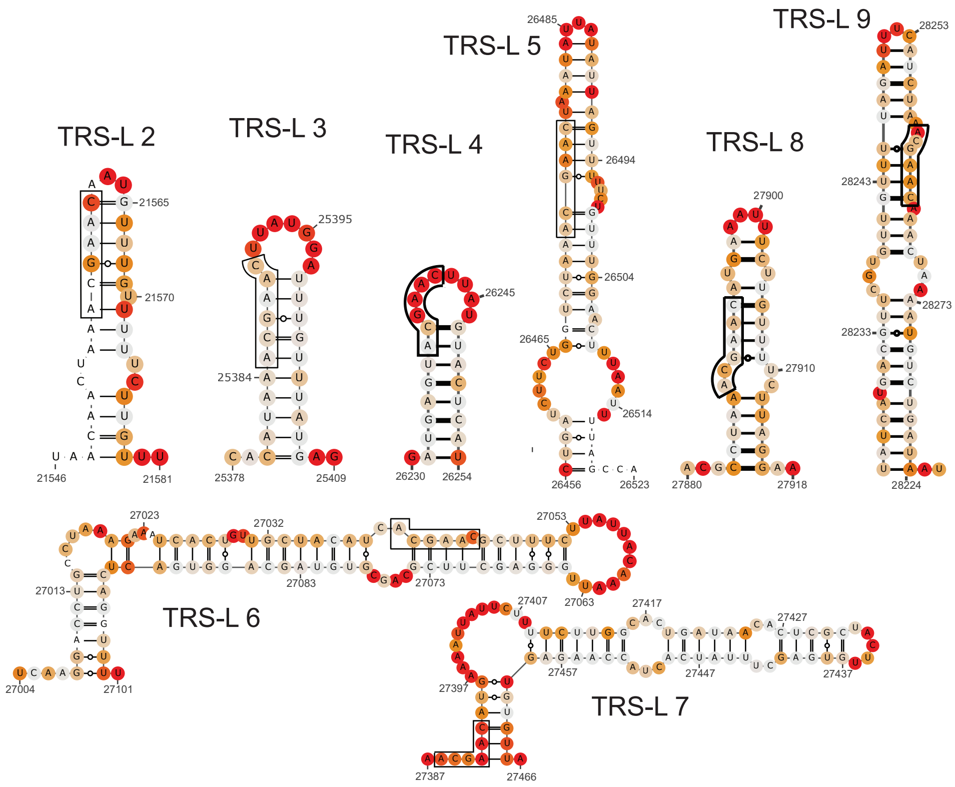


**c**

| | WT | Delta382 | DEN1 | ZILM |
|---|---|---|---|---|
| Base Count | 29844 | 29463 | 10735 | 10807 |
| Paired Bases | 17060 | 16968 | 5396 | 5976 |
| Paired Bases (%) | 57.16 | 57.59 | 50.27 | 55.3 |
| Median Base Pair Span (nt) | 26 | 26 | 35 | 32 |
| Average Base Pair Span (nt) | 60.76 | 64.05 | 84.4 | 75.05 |
| Median Helix Length (nt) | 5 | 5 | 4 | 5 |
| Average Helix Length (nt) | 5.54 | 5.54 | 5.06 | 5.48 |
| Max Helix Length (nt) | 27 | 28 | 24 | 27 |
| Min Helix Length (nt) | 1 | 1 | 1 | 1 |
